# scConcept: Contrastive pretraining for technology-agnostic single-cell representations beyond reconstruction

**DOI:** 10.1101/2025.10.14.682419

**Authors:** Mojtaba Bahrami, Alejandro Tejada-Lapuerta, Sören Becker, Fatemeh S. Hashemi G., Fabian J. Theis

**Affiliations:** Institute of Computational Biology, Computational Health Center, Helmholtz Munich, Germany; School of Life Sciences Weihenstephan, Technical University of Munich, Germany; School of Computation, Information and Technology, Technical University of Munich, Germany; Munich Center for Machine Learning, Ludwig-Maximilians-Universität München, Germany

## Abstract

Recent large-scale single-cell foundation models have shown promise for exploring cellular states, yet they often underperform compared to simpler, domain-specific methods, raising concerns about their broader applicability. A key limitation lies in their reliance on masked language modeling, which is well suited for generative language tasks but poorly aligned with learning rich cell-level embeddings required in single-cell biology. Moreover, the proliferation of transcriptomic technologies—from whole transcriptome dissociated assays to image-based targeted profiling—poses a major challenge for cross-platform generalization. Here, we align with recent advances in machine learning to move beyond reconstruction metrics, which often do not capture important sample variation. We present scConcept (“contrastive cell pre-training”), a transformer-based contrastive learning framework that directly optimizes cell embeddings by contrasting multiple views of cells. By replacing gene-level reconstruction with a cell-level identification task, scConcept learns robust representations that are invariant to count distributions and gene panel selection, across diverse assays and technologies. To highlight the capability of the proposed framework, we pretrain scConcept on a similar corpus of over 30 million single-cell RNA-seq profiles as recent foundation models. Our approach demonstrates superior performance not only compared to state-of-the-art pretrained foundation models but also domain-specific methods in various downstream tasks, including cell-type annotation, technology integration, dissociated to spatial cell-type transfer, spatial imputation, gene panel optimization, and mapping new technologies on already existing atlases. Our results highlight contrastive pretraining as a powerful alternative to reconstruction-based strategies for single-cell modeling, providing a path toward general-purpose, technology-agnostic cell representations.

## Introduction

Recent advances in the cost-effective execution of single-cell transcriptomics have led to the accumulation of large-scale scRNA-seq datasets across diverse tissues. Motivated by the success of self-supervised pre-training in large language models ^1–4^, recent efforts ^5–11^ have focused on developing analogous large-scale models for single-cell omics, commonly referred to as foundation models. While Masked Language Modeling (MLM) and Next Token Prediction (NTP) (also known as causal language modeling) have shown to be effective pretraining tasks in language understanding ^2,12,13^ and generative language modeling ^1,14,15^, adopting a similar gene-level masking approach to model cellular states still struggles to outperform simple or domain-specific models on most downstream tasks ^16–19^.

This discrepancy is partially caused by the fact that the objective of a generative natural language model is consistent across training and inference time, which is predicting the next token in a sequence. Therefore, as long as the input sequences are not out of distribution, the model is evaluated through solving the same task for which it is trained for–without any concern over the underlying learned representations and their semantics. In contrast, the pre-training objectives of cell foundation models mostly serve as a proxy task, where the latent representations of the model are considered as the main asset of it which can be used for exploratory analysis or as rich features for solving downstream tasks. Despite this key difference in the use case, almost all the recent single-cell foundation models ^5–11^ use gene-level masked language modeling without introducing a solid objective for directly learning cell-level representations which are used as main products of such models. To generate cell embeddings, most single-cell foundation models compute a uniform or weighted average of contextualized gene embeddings. However, these uninformed averages do not necessarily yield optimal cellular representations, which can limit their effectiveness on downstream tasks. As a result, they may struggle in scenarios where simpler or domain-specific models—such as PCA, linear approaches, or VAE-based methods—explicitly learn cell embeddings or solve supervised tasks ^20–24^. On the other hand, unlike language generative models, natural language understanding use cases that care about the learned representations have tried to introduce secondary objectives, such as “*next sentence prediction”* ^2^ and “*sentence ordering*” ^25^ to learn high-level sentence semantics beyond words. However, the distinct nature of single-cell omics data—lacking elements such as sentence relationships—limits the direct application of such approaches.

Moreover, the diverse range of transcriptomic technologies and assays from whole transcriptome ^26–30^ to in-situ microscopic methods ^31–33^ has posed a substantial challenge for building technology-independent cell foundation models that are able to learn and extract the shared biological phenomena regardless of technology biases and differences, including differences in count distributions and target gene panels. Training on different technologies ^8,34^ has led to limited improvement on unseen data since it has not been directly addressed in the model design and training to improve transferability to different and even unseen technologies.

Contrastive learning is a powerful self-supervised framework that has achieved remarkable success in both computer vision and natural language processing domains ^35–41^. Several studies have explored contrastive learning in the context of single-cell transcriptomics ^42–50^. However, these works have either focused on using contrastive loss for data modality integration or were not intended for large-scale model pre-training. A central challenge in applying contrastive learning to transcriptomic data lies in constructing meaningful positive augmented pairs and determining appropriate negative samples. Existing approaches have relied on combinations of expression-level noise, cell type–based augmentations, and gene dropout strategies. Nevertheless, recent evidence ^51^ suggests that with such limited augmentations, contrastive learning approaches remain suboptimal and exhibit diminished performance.

In this work, we propose scConcept, a transformer-based contrastive learning framework for large-scale self-supervised pretraining on transcriptomic data. scConcept learns robust and expressive cell representations that generalize across varying count distributions and gene panels from diverse assays and technologies. By replacing gene-level reconstruction with a more effective cell-level identification task, our approach directly optimizes for the cell embeddings that are used in downstream single-cell analyses. Here we highlight the effectiveness of the proposed architecture by limiting the pre-training data to only dissociated single-cell RNA-seq data with the same scale and diversity as the recent masked language modeling foundation models of scGPT and Geneformer. We also compare with Nicheformer, which is pretrained over a larger corpus of both single-cell and spatial transcriptomics. We further show that the unique architectural design of scConcept facilitates natural generalization to different spatial technologies while only being pre-trained over dissociated scRNA-seq profiles.

## Results

### A contrastive transformer-based model for learning robust cell representations

scConcept is an alternative approach to masked language modeling (MLM) for pre-training cellular representations through contrastive learning. Contrastive learning is a powerful self-supervised learning paradigm to learn by comparing positive and negative sample pairs. It enables the model to learn meaningful representations by bringing relevant data points closer and pushing irrelevant ones apart in the embedding space, resulting in a rich representation space fulfilling desired properties of interest. We show that contrasting cell identities (or cell views) is a superior alternative pre-training task that is capable of learning rich and robust cell representations usable in different downstream tasks. scConcept is a standard transformer-based architecture consisting of stacked blocks of multi-head self-attention followed by feed-forward networks and layer normalizations connected through skip connections as specified in the original work ^14^.

scConcept is trained to learn representations of cell views, which we define as any arbitrary subset of genes from a cell’s expression profile. Cell views can be either a random or an informed subset of genes, like pre-designed gene panels used in targeted spatial transcriptomics. scConcept is optimized to produce similar representations for a disjoint pair of views originating from the same cell while maximizing their distance to views of other cells through contrastive learning (Fig. 1a-b). Training the model to obtain consistent cell embeddings regardless of the subset of genes in the gene panel encourages robustness and generalization, enabling scConcept to capture the underlying cellular identity despite variability in gene panels. One of the main advantages of contrasting cells or cell views with one another is that it inherently facilitates the learning of cell-level representations—an outcome that is not directly achieved by reconstructing masked genes in the model’s output. To enable this, a [CLS] token is appended to the input sequence of each cell view, and its corresponding output embedding is used to represent the entire cell view (Fig. 1c-d). A similarity matrix is then constructed by computing the dot products between all pairs of embeddings. These embeddings are contrasted using the proposed modified InfoNCE loss ^35^, as detailed in Methods.

**Figure 1:**
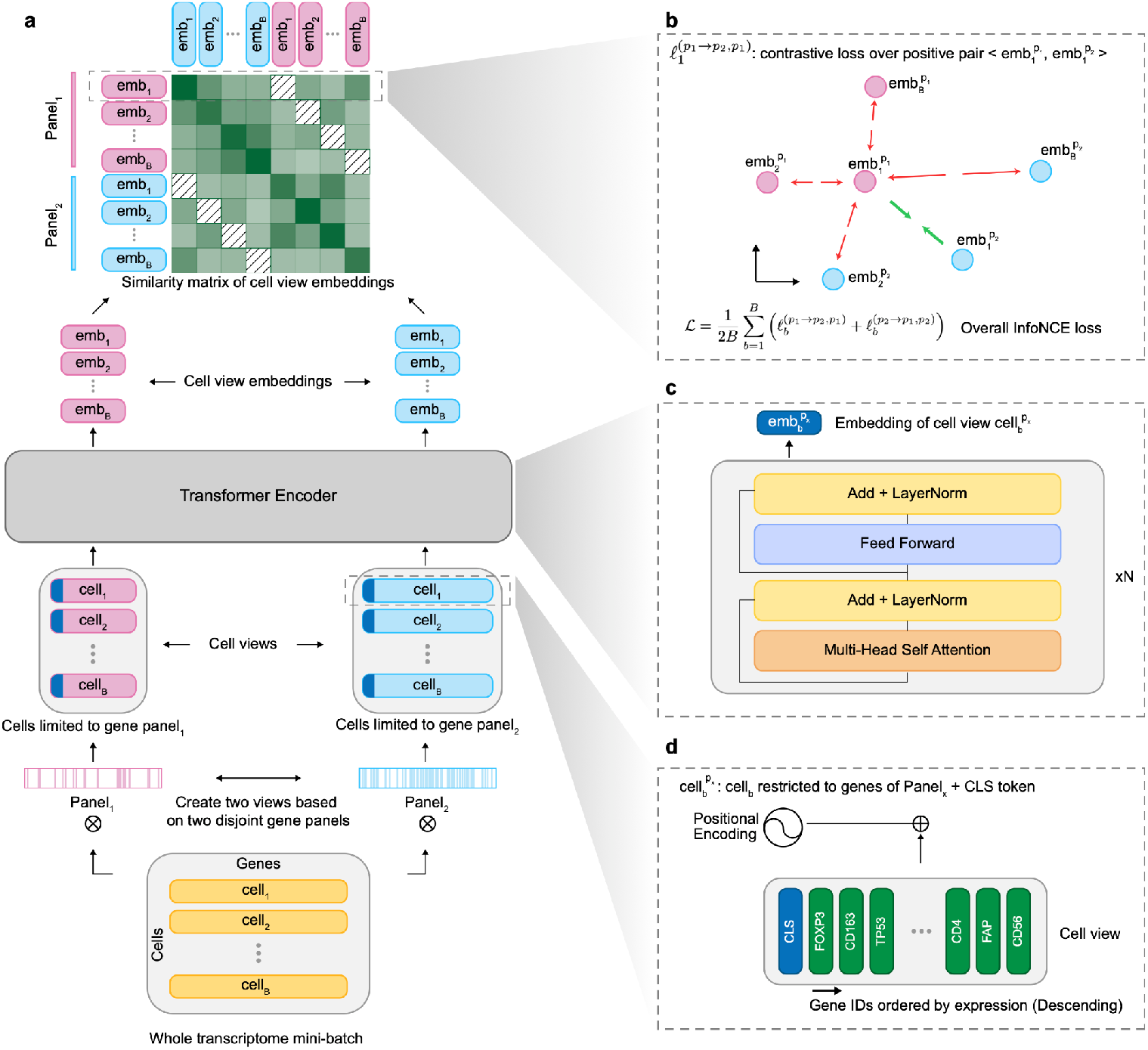
scConcept model and training overview. (**a**) scConcept is trained to produce consistent cell embeddings for any subset of selected genes known as gene panels. At each training step, a pair of disjoint gene panels is sampled from the pool of random or pre-designed gene panels. Each cell in the mini-batch is then subsetted to genes in each panel independently and creating two views of the same cell. A transformer encoder architecture is then trained to produce corresponding cell view embeddings in a way that optimizes the output similarity matrix of all cell views through InfoNCE contrastive loss. (**b**) The contrastive objective encourages the network to produce similar cell embeddings for the cell views originating from the same cell (the positive pair) and dissimilar embeddings for the pairs originating from different cells (negative pairs). Each cell is not only contrasted against the cells from the opposite view but also with all the instances from the same view, which contain the same gene panel as the cell of interest. (**c-d**) Every cell in each view is first rank encoded by sorting its genes in a descending order based on the absolute expression values (**d**), and then is processed through a stack of Nx Transformer Blocks from which the output embedding of the [CLS] token is selected as the representing embedding of the corresponding cell view (**c**).

We identify two additional key modifications that considerably enhance the learning of richer and more robust cell embeddings. First, to ensure homogeneous embeddings that are independent of specific gene panels, we contrast each cell not only with cells from other views but also with the remaining cells within its own native view. This strategy promotes the dominance of positive pair similarity over intra-view similarity. In other words, we aim for a cell’s embedding to closely align with its positive counterpart from the opposite view, while also being distinguishable from other cells within the same view. This dual objective ensures that the learned representations capture unique, discriminative features that go beyond mere cross-view alignment. Second, rather than randomly sampling cells from the entire pre-training corpus, we find it more effective to construct each mini-batch using cells drawn exclusively from a single dataset. The underlying intuition is that distinguishing between cells of the same or closely related cell types within a dataset (e.g., distinguishing immune cell subtypes from PBMC dataset) forces the model to learn finer-grained, biologically relevant features. In contrast, separating clearly distinct types (e.g., immune vs. neural cells from different studies) is comparatively easy and offers less signal for learning subtle representations. By focusing comparisons within a dataset, we also avoid driving the model to use inter-dataset technical differences—rather than biology—as distinguishing features, thereby mitigating batch effects and improving the relevance of the learned representations.

For any given hold-out dataset, the quality and performance of a pre-trained model’s embeddings largely depend on the similarity between the new data and the data encountered during pre-training. Ideally, a foundation model trained on a sufficiently large and diverse corpus—including data from a wide range of tissues and organs—would generalize well to more hold-out datasets. However, in the absence of such comprehensive pre-training data, distribution mismatches between new target datasets and the pre-training data are inevitable. These discrepancies can be mitigated by adapting the pre-trained model to the target data at inference time. A straightforward strategy for adaptation is to continue the self-supervised pre-training of the model on the new target data for a few additional steps. We refer to the resulting model as scConcept+. Unlike task-specific fine-tuning, this form of model adaptation is performed in a task-independent manner, without the use of any labels or supervision. In contrast, methods that have not been adapted to the evaluation dataset are commonly referred to as zero-shot methods. We evaluate the performance of scConcept against recently proposed zero-shot methods, including scGPT, Geneformer, and Nicheformer. In addition, we benchmark the domain-adapted variant, scConcept+, against PCA, scVI, Tangram, Celltypist, and scArches across different downstream tasks optimized on the evaluation dataset.

### scConcept enables accurate cell-type annotation across tissues

Cell-type annotation is a critical step in understanding the cellular composition and function of complex tissues, enabling insights into development, disease mechanisms, and therapeutic targets. To evaluate the performance of the scConcept embeddings for the task of cell-type annotation, we take three scRNA-seq datasets from different tissues and split each into a pair of reference and query sets based on donors, measurement site, and disease state respectively. We then extract the embeddings of the model for both reference and query sets. The query set is then annotated using a KNN classifier by finding the nearest neighbors of each query cell from the reference set through majority voting (Methods). Given the predicted cell types, we then measure the accuracy and macro F1 score to evaluate the performance of the predicted annotations. We compare the results to the predictions using the embeddings extracted from scGPT, Geneformer, and Nicheformer. We also include PCA and CellTypist^52^ as domain-specific methods for comparison (Methods).

First, we take a bone marrow dataset of 42,492 mononuclear cells from 12 healthy human donors collected across three separate experiment sites ^53^. We take the data from the first site as the reference split and predict the annotations for the data from the second site, simulating a real scenario where query and reference datasets are sampled and sequenced separately in different conditions. The dataset comprises 22 cell types with an uneven distribution across the different classes (Fig. 2d) for both reference and query datasets. According to the original study, the cells are annotated using the joint transcriptome and chromatin accessibility profiles together as the data is part of a multiome assay, however here we only exploit the gene expression modality for annotation. The overall performance of scConcept (Fig. 2a) shows at least 5% improvement of macro F1 score and 4% improvement of accuracy compared to other foundation models evaluated in the zero-shot setting. It also outperforms simple and domain-specific baselines (i.e. PCA and CellTypist) on both accuracy and macro F1 score. By adapting the pre-trained model over the target dataset, the model performance (scConcept+) can be boosted by an additional 1.8% in macro F1 score (Methods).

**Figure 2:**
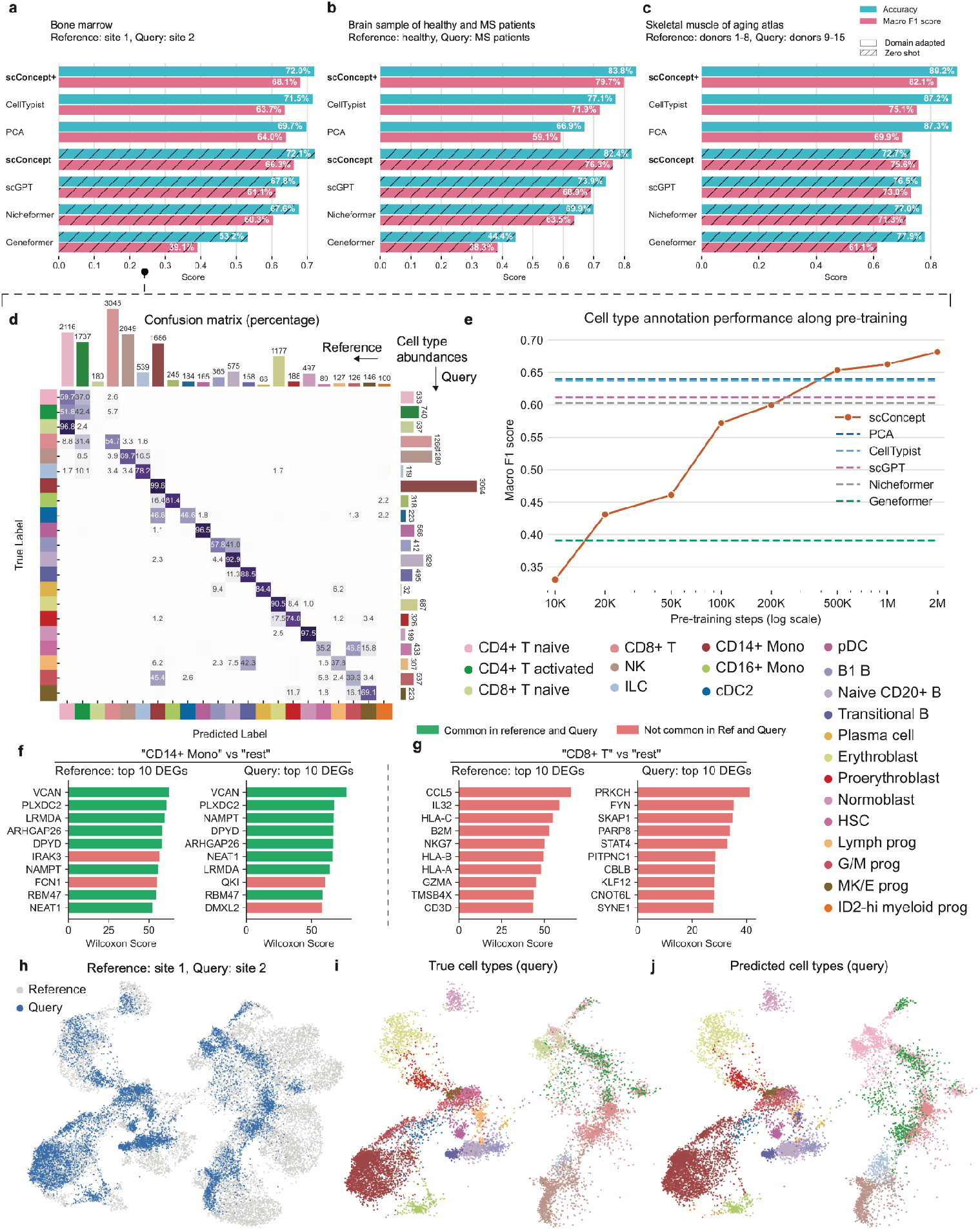
scConcept learns high-performing cell representations, boosting cell type identification. (**a–c**) Accuracy and macro F1 scores for cell-type annotation comparing different methods on three datasets: bone marrow across different experimental sites (**a**), brain from healthy and MS donors (**b**), and skeletal muscle across donors (**c**). The proposed method (scConcept) and the adapted version (scConcept+) outperform masked language modeling-based foundation models (scGPT, Geneformer, and Nicheformer) and domain-specific baselines (PCA, CellTypist), respectively. (**d**) Confusion matrix showing predicted vs. true cell types in the bone marrow dataset using scConcept embeddings, with most misannotations occurring between closely related cell types. (**e**) Macro F1 score of scConcept at various pre-training steps, demonstrating continued performance gains along the pre-training. (**f-g**) The top DEGs of CD14+ monocytes and CD8+ T cells, calculated separately for reference and query sets. DEGs in common are in green, otherwise in red. (**h–j**) UMAP visualizations of embeddings from the bone marrow dataset: reference and query sites (**h**), true cell-type labels (**i**), and predicted cell-type labels (**j**). Despite the existing batch effect between the sites, scConcept embeddings enable accurate label transfer across datasets.

Looking into the details of the model’s predictions (Fig. 2d), most instances of apparent misannotation occur between closely related cell types or maturation states within the same lineage (e.g., B cell subtypes or progenitor-to-mature transitions), reflecting biological continuity rather than major classification errors. However, a major part of the mis-annotation is caused by the inconsistency of a cell-type signature across the reference and query sets which could be due to the batch effect between the two sites of measurements. For example, we observe (Fig. 2f) that for *CD14+ monocytes*, many of the top 10 differentially expressed genes (DEGs) are shared between the reference and query sets and consequently the model has a very good performance in identifying them. However, for *CD8+ T cells*, there is no gene in common between the top 10 DEGs of the reference and query sets (Fig. 2g) which partly explains why they are not well identified by scConcept as they manifest different signatures. A similar relation is also visible for other cell types (Supplementary Fig. S3).

We also plot the UMAP of the model embeddings (Fig. 2h) for reference and query samples. As the reference and query splits come from different experiment sites, a small drift is seen showing some batch effect between different sites. However, despite the shift, we see that scConcept is able to embed the semantically related cell types close to each other (Fig. 2i-j).

To illustrate the effect of the pre-training stage in learning high-performance representations, we plot the performance (macro F1 score) of the model at different training checkpoints through the pre-training (Fig. 2e). The model outperforms all other models after 500K training steps and keeps improving further as training proceeds. Although the performance trend suggests that further training leads to further performance improvement, these improvements come at higher computational cost (as indicated by the logarithmic scale of the x-axis in Fig. 2e), which is why we choose the 1M checkpoint for all subsequent analyses.

To assess the generalization ability of the learned embedding space across different physiological and disease states, we try a second dataset that consists of brain samples from 9 healthy controls and 12 individuals with Multiple Sclerosis (MS) ^54^. Accordingly, we partition the data into a reference set of healthy donors and a query set of MS patients. Consistent with previous observations, scConcept demonstrates strong generalization by accurately capturing cell identities across varying conditions (Fig. 2b).

To further evaluate the model’s data efficiency, we incorporate a less-represented tissue type during pre-training. Specifically, we use the human skeletal muscle aging atlas dataset ^55^, which includes samples from 15 donors spanning the adult human lifespan. We designate half of the donors (1–8) as the reference set and the remaining (9–15) as query cells. Despite skeletal muscle tissue being underrepresented in the pre-training data, scConcept performs comparably well (Fig. 2c), demonstrating a more data-efficient pre-training scheme. Notably, the adapted model (scConcept+) achieves a substantial performance boost, highlighting the potential for improvement through the inclusion of more data from the target tissue in the pre-training phase.

### scConcept facilitates generalization to spatial transcriptomics

Recent advances in spatial transcriptomics have enabled the high-resolution mapping of cellular architecture, providing unprecedented insights into cellular composition and subcellular interactions. However, the inherent technological biases and the constrained gene panels of imaging-based methods limit the direct translatability of insights derived from comprehensive, whole-transcriptome scRNA-seq datasets. A key challenge lies in accurately transferring well-curated cell-type annotations from scRNA-seq references to novel spatial transcriptomic data. The success of this label transfer critically depends on the fidelity and alignment of the shared representation space between the two readouts. Here we demonstrate the natural generalization of scConcept, which has only been pre-trained on dissociated scRNA-seq data to unseen spatial technologies, showing the effectiveness of the proposed pre-training scheme. For this purpose, we first perform a simulated experiment to understand the dependency of the learned embeddings on the input gene panel, followed by two real world scenarios of transferring cell-type labels of pairs of real dissociated and spatial technologies of the same tissue and condition.

We take a bone marrow dataset of mononuclear cells from 12 healthy human donors across 3 different experiment sites. The dataset contains the count values mapped to 13,431 genes. We create a pair of reference and query subsets based on the corresponding experiment sites, resulting in a reference subset of donors 1-3 (from site 1) and a query subset of donors 4-7 (from site 2). We take a subset of 265 genes based on a pre-designed Immuno-Oncology Xenium panel from 10x Genomics and create a pseudo-spatial query set (Fig. 3a). We then extract the embeddings from scConcept for both the reference and query sets respectively. We show the UMAP ^56^ visualizations of the output cell embeddings of scConcept along with scGPT, Geneformer, and Nicheformer for the pair of whole transcriptome scRNA-seq reference and the pseudo spatial query in Fig. 3b. We clearly see that scConcept is able to learn gene panel independent cell representations, enabling integrative analysis of different technologies with different gene panels. We then measure the cell type transfer performance by training a KNN classifier for cell-type annotation over the reference set and evaluate its performance over both the full transcriptome and pseudo Xenium panel query sets. As observed in Fig. 3c, the performance (macro F1 score) of the limited Xenium panel expectedly drops compared to the full transcriptome case. This performance drop can partly be attributed to the inherently limited information content in the subset of genes in the Xenium panel, but is in addition also caused by the natural sensitivity of the neural architectures including transformers against the number of input genes and therefore input sequence length which impacts the output representation. We observe that scConcept shows the least performance drop, which clearly indicates its superior robustness with respect to the input gene panel. As the number of features (genes) are different across the two reference and query sets, a feature standardization is required for the comparison to non-transformer-based baselines such as PCA (Methods). We further looked into the mis-annotations of the scConcept (Supplementary Fig. S4), causing the performance drop when switching to the limited Xenium panel query set, and measured the availability of the key cell type markers in the pseudo Xenium panel. For this, we run a differential gene expression analysis using the Wilcoxon test for each cell type group compared to the rest of the cells and assess which and how many of the top differentially expressed genes (DEGs) are present in the reduced Xenium panel (Fig. 3d). We see that for cell types with good cell type transfer performance (e.g. natural killer cells), a sufficient number of markers are still present in the limited Xenium panel; however, the cell types with the highest performance drop (e.g. Normoblast and cDC2 dendritic cells) are the ones with very few or no markers left in the Xenium panel (refer to supplementary Fig. S5 for all cell types). An overall similar trend can also be seen by comparing the celltype annotation performance with respect to the rate of the top 30 DEGs of each cell-type present in the Xenium panel (Fig. 3e).

**Figure 3:**
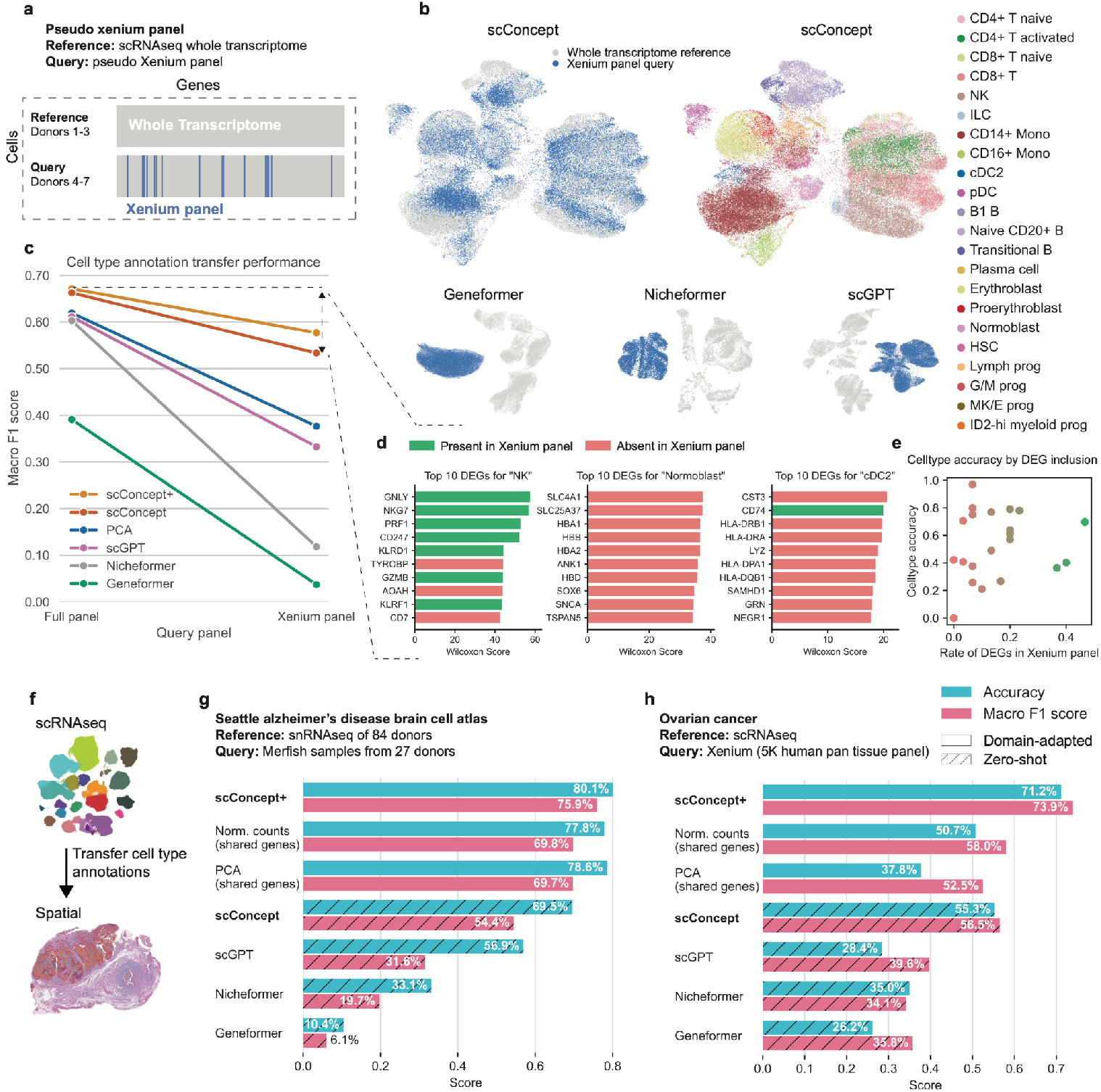
scConcept enables robust label transfer across technologies. (**a-d**) Experiment of gene panel independence. (**a**) The experiment setup in which a whole transcriptome scRNA-seq dataset of bone marrow samples is separated into reference and query sets based on the experiment sites, where the query set is subsetted to a pre-defined xenium panel. (**b**) UMAP plots of the co-embedding of the whole transcriptome and the Xenium Immuno-Oncology panel of the same scRNA-seq dataset, showing the robustness of the scConcept pre-training to different gene panels. (**c**) The robustness of cell embeddings for cell-type transfer against different query gene panels. (**d**) The presence of the cell type signatures (top 10 DEGs) of example cell types in the pseudo Xenium panel. The genes are colored in green if they are present in the pseudo Xenium panel subset, otherwise in red. (**e**) The celltype accuracies based on the rate of top 30 DEGs present in the Xenium panel. (**f-h**) Cell-type transfer accuracy between real dissociated scRNA-seq and targeted spatial transcriptomic. (**f**) The experiment setup of co-embedding a scRNA-seq (reference) and targeted spatial transcriptomic (query) and transferring the cell-type labels. (**g**) The transferability performance of Alzheimer’s brain samples from scRNA-seq to MERFISH samples through training a linear head on the scRNA-seq reference. (**h**) The same performance for annotating a Xenium 5K Prime sample using a scRNA-seq reference of ovarian cancer.

Next, we evaluate the capability of cell-type transfer between pairs of real whole transcriptome and image-based targeted spatial transcriptomic datasets (Fig. 3f). First, we take the Seattle Alzheimer’s Disease Brain Cell Atlas ^57^ that includes single-nucleus RNA sequencing (snRNA-seq) and cellularly resolved multiplexed error-robust fluorescence in situ hybridization (MERFISH) of the middle temporal gyrus (MTG) from a cohort of 84 and 27 aged donors respectively, spanning the spectrum of AD pathology. The snRNA-seq and the MERFISH readouts contain the expression profile of 36,412 and 139 genes, respectively, that expand across 21 cell types. We embed all the cells from both snRNA-seq and MERFISH using scConcept and train a linear classifier over the embeddings of the snRNA-seq subset as a reference and evaluate the cell type annotations on the MERFISH query set in terms of accuracy and macro F1 score. The cell representations obtained from scConcept enable a strong cell-type label transfer from snRNA-seq to the MERFISH technology compared to other foundation models (Fig. 3g). We also include normalized counts and PCA embeddings as baselines in our comparison. Each modality is subset to the 139 shared genes, and we use either the normalized counts directly or their PCA-reduced representations for label transfer. The strong performance of the baseline methods can be partly explained by their similarity to the original annotation procedure, which relies on comparing normalized counts of shared genes between the query and reference modalities. Although scConcept achieves the best performance among zero-shot methods, it only slightly outperforms using normalized counts alone. This limited improvement can be attributed to two factors: the biased evaluation setup, which favors methods resembling the original annotation procedure, and the restricted diversity of the pre-training dataset. These constraints on zero-shot performance can be mitigated by domain-specific fine-tuning, as demonstrated by the performance of scConcept+, which addresses these issues and consistently outperforms all other methods.

Finally we assess cell type label transfer from samples of ovarian cancer from a scRNA-seq reference to the query dataset of the Xenium assay of human ovarian cancer with the 5K Prime human pan tissue Xenium panel from 10x genomics (Fig. 3h). The scRNA-seq and the 5K Prime Xenium panel contain the measurement of 18,082 and 5,101 genes, respectively, with 4,912 genes in common between them. Both datasets are annotated with 16 harmonized cell type labels that enable evaluation of the cell type transfer performance. In this case, scConcept shows a comparative performance as the PCA and normalized count baselines without any further adaptation and outperforms the other foundation model approaches with a strong margin. Adapting scConcept+ on the reference and query datasets yields a macro F1 of 0.739 and accuracy of 0.712, clearly surpassing all baselines.

### scConcept enables reference-based spatial transcriptomic imputation

A diverse array of spatial transcriptomic technologies exists, each presenting unique limitations regarding cellular resolution and the breadth of the gene panel they can capture. High-resolution imaging-based technologies, such as seqFISH ^33^, MERFISH ^32^, and commercial platforms like Xenium, MERSCOPE, and CosMx, can pinpoint individual RNA molecules with subcellular precision. However, these technologies are currently constrained to measuring a limited number of genes in a single experiment due to inherent technological restrictions. Consequently, transcriptomic imputation becomes a vital step for achieving a comprehensive understanding of underlying biological processes. Most imputation methods leverage corresponding whole-transcriptome single-cell RNA sequencing (scRNA-seq) reference datasets to predict the expression of unobserved genes in spatial datasets. Ideally, these reference datasets should originate from a biologically similar system, if not the same sample, to ensure accurate imputation.

Unlike cell type label transfer from whole-transcriptome references to spatial assays—where labels are often derived through manual annotation—gene imputation focuses on predicting normalized expression counts, the primary and unprocessed outputs of the experiments. This provides a more direct and unbiased basis for evaluating cell representations. Such evaluation is less susceptible to errors introduced during the annotation process and avoids circular dependencies that can arise when PCA and shared gene subsetting strategies are used through the labeling workflow. Conversely, accurate gene expression imputation may prove more difficult due to the inherent measurement noise in the data, a challenge which is further compounded by the fact that noise distributions associated with distinct technologies may differ.

To evaluate gene imputation in real spatial assays where the true expression of unmeasured genes is unknown, we hold out a subset of already measured genes. These held-out genes are then used to assess the performance of the imputation predictions. Since there are always a number of genes with very low or zero variation in the spatial panel, we evaluate the performance of imputation over the subset of highly variable genes of the hold-out gene set (Fig. 4a). We impute the hold-out genes by first extracting embeddings from the spatial query and the whole transcriptome reference, then identifying the k=10 nearest neighbors for each spatial cell among the reference cells, and finally predicting the imputed expression as the average hold-out gene expression of these neighbors. Subsequently, the Pearson correlation coefficient (PCC) is computed for each gene independently across all cells, comparing true and predicted expression counts. The average of these gene-specific PCCs then serves as an overall performance score for each method (Methods).

**Figure 4:**
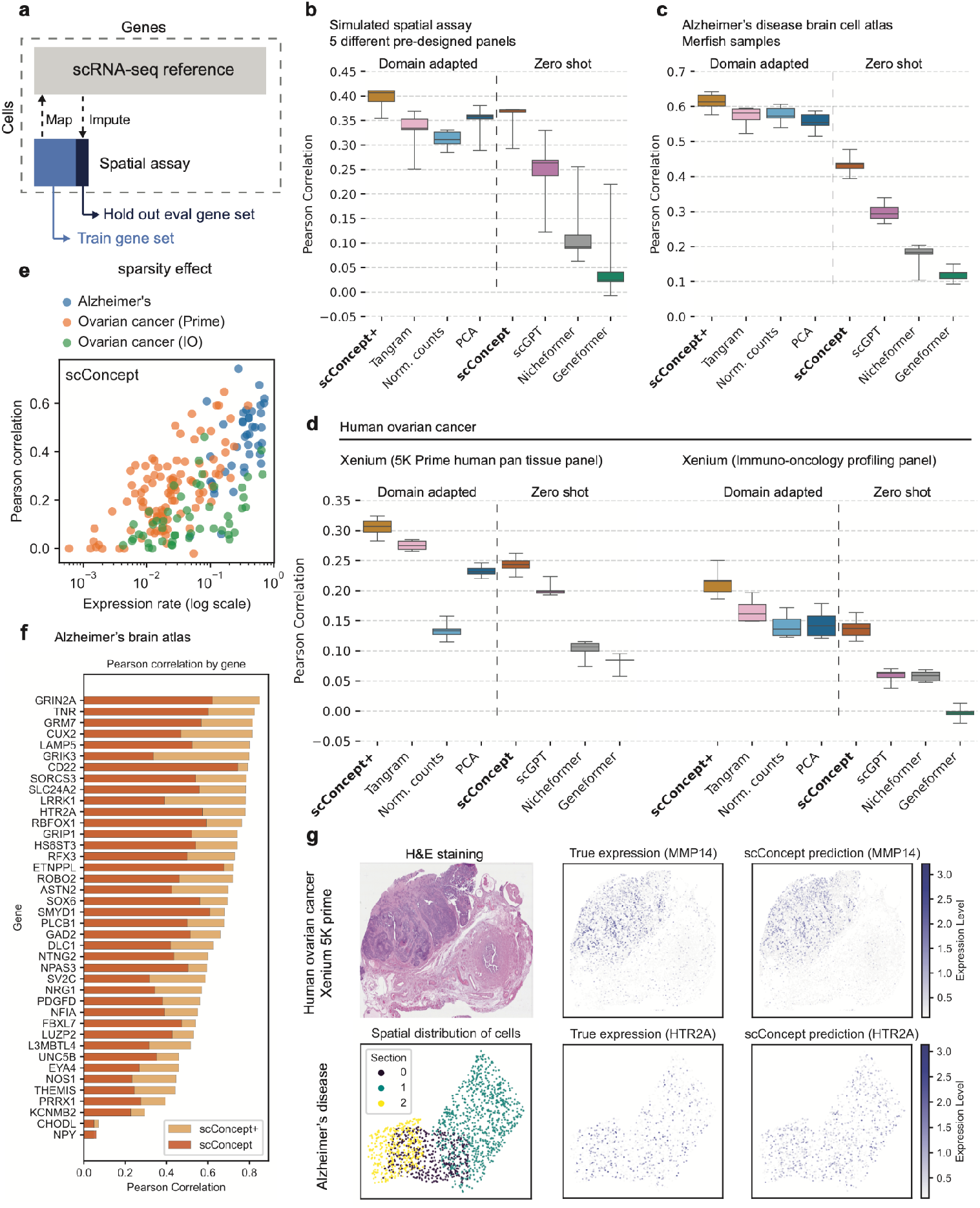
scConcept enables accurate gene imputation in targeted spatial assays. (**a**) The schematic of the imputation task evaluation where a portion of the spatial genes is held out for evaluating the imputation performance. (**b-d**) The Pearson Correlation Coefficient (PCC) of the gene imputation task compared with the performance of the embeddings derived from other foundation models and the baselines in the two zero-shot and domain-adapted settings, respectively. We impute the top highly variable genes (HVGs) for a scRNA-seq pseudo xenium panel query set in (b), the top HVGs of a MERFISH Alzheimer’s Disease brain dataset in (c), and the top HVGs of a 5K Human Pan Tissue Xenium and an Immuno-Oncology Xenium data, both from human ovarian cancer datasets in (d). (**e**) The effect of gene capture efficiency of different genes from three different assays on the imputation performance for different datasets. (**f**) The imputation performance gain obtained by the adapted version of the model (scConcept+) across the 40 held-out genes in the MERFISH Alzheimer’s disease data. (**g**) The example spatial distribution of the true expressions and the imputed values for the gene MMP14 of the ovarian cancer dataset and HTR2A of the MERFISH Alzheimer’s Disease brain dataset, where both are known to have important regulatory roles in Alzheimer’s and ovarian cancer, respectively. scConcept is able to recover the spatial variation of the corresponding genes.

Any distribution mismatch due to batch effect between the whole transcriptome reference and the limited panel spatial assay can affect the performance of the imputation and distort the method evaluation. To have a batch effect-free evaluation, we first try a pseudo-experiment where we take a bone marrow whole transcriptome scRNA-seq dataset and create a pair of whole transcriptome reference and a pseudo spatial query set by subsetting the whole transcriptome data to the genes available in a few existing panels of interest (Methods). We compare the overall PCC score of scConcept and scConcept+ with the results from the embeddings obtained by other foundation models. We also compare with the k-nearest neighbor approach over the normalized and log1p-transformed count space (Norm. count), PCA space, and also the domain-specific method of Tangram ^58^ (Methods), which was shown to be among the best-performing currently available domain-specific imputation methods ^59^. scConcept shows a superior PCC compared to other methods (Fig. 4b), surpassing the performance of not only other foundation models in the zero-shot setting, but also the domain-specific approaches. Additional domain adaptation further boosts this performance as evidenced in scConcept+.

We then proceed to evaluate the imputation performance on actual spatial technologies. For this, we take the MERFISH assay of 139 targeted genes from 27 donors of the Alzheimer’s Disease Brain Cell Atlas ^57^ dataset with a corresponding snRNA-seq reference from 84 donors. The snRNA-seq reference contains 36,412 genes including all 139 genes of the spatial assay. We run a 6-fold evaluation, where we hold out a set of 40 random genes and use the remaining 99 genes for training and prediction at each iteration. As demonstrated in Fig. 4c scConcept shows a strong performance in the zero-shot setting compared to other foundation models, however does not perform as well as domain adapted methods (i.e. Normalized counts, PCA, Tangram). However, the scConcept+ is able to get the best performance after being adapted over the target pairs of reference and query datasets. We further investigated a distinct dataset derived from 10x assays of an ovarian cancer sample, comprising a 10x scRNA-seq whole transcriptome sample alongside two 10x Xenium spatial assays: the 5K human pan tissue panel and the immuno-oncology profiling panel. A similar trend is evident in both of these spatial datasets, respectively (Fig. 4d), thereby demonstrating the robust and generalizable cell representations obtained through scConcept across different technologies and gene panels. We also compare the methods by Spearman’s rank correlation coefficient (Supplementary Fig. S6) and observe a similar trend and superior performance of scConcept. While Pearson’s correlation assesses linear relationships, Spearman’s rank correlation assesses monotonic relationships whether linear or not.

The imputation performance in the three spatial datasets relies on the different factors, including the quality of the spatial assay. By looking into the effect of the sparsity of different genes in different datasets on the imputation performance (Fig. 4e), we observe a trend showing a generally better imputation performance for genes that are expressed in more cells, which implicitly relates the data quality to the overall imputation performance. In our case, the MERFISH assay in Alzheimer’s brain atlas shows a higher transcript capture efficiency resulting in better imputation performance in general. We also examine the imputation performance for each individual gene in Fig. 4f for one of the gene splits and show the performance gain achieved by scConcept+ through the unsupervised adaptation over the target datasets. We see that scConcept+ consistently improves the imputation performance in all hold-out genes in addition to the net increase in the overall performance without any negative impact as a trade-off. Moreover, the magnitude of improvement varies across genes, rather than manifesting as a uniform gain for all.

Looking into the spatial slides (Fig. 4g) of the imputed expression counts against the true expression counts for *MMP14*, which plays a significant role in human ovarian cancer, its prognosis, progression, and metastasis ^60,61^ shows the relatively strong capability of scConcept to reconstruct the spatial expression pattern of this gene. A similar pattern is observed for the *HTR2A* gene, which has been shown to play a role in Alzheimer’s disease, particularly in relation to associated symptoms like psychosis and depression ^62^ and episodic memory consolidation ^63^. By mapping the imputed expressions of *MMP14* and *HTR2A* onto the native spatial organization of their corresponding cells, we observe that scConcept effectively recovers the spatial variation of these previously unobserved genes of interest compared to other methods (Supplementary Fig. S7).

### scConcept facilitates panel design and optimization

Most single-cell data analysis pipelines focus on single gene statistics, including highly variable genes (HVGs) and differentially expressed genes (DEGs), or pairwise gene correlation metrics that capture the linear or rank-based co-expression patterns of pairs of single genes. Likewise, most unsupervised approaches used for designing optimal gene panels for targeted spatial assays that maximally capture signal variation either rely on single-gene statistics or optimize for indirect gene-set metrics and heuristics ^64–69^. On the other hand, mutual information (MI) is a suitable statistical measure for quantifying the information shared between different sets of random variables (e.g., gene sets). However, in practice, we rarely know the true joint probability distribution needed to compute MI directly. To address this limitation, contrastive frameworks can estimate mutual information from limited samples. Here, we propose scConcept as a framework for measuring the information dependency between any two arbitrary sets of genes within a given dataset (Methods).

To estimate the MI between any two gene panels using scConcept+, a mini-batch of cells is required. We note that the size of this batch strongly influences estimation accuracy. In Fig. 5a, we show the MI between a Xenium immuno-oncology panel and the entire transcriptome, computed on a bone marrow scRNA-seq dataset. As the number of cells increases, the MI estimate stabilizes, with convergence observed at around 2,048 cells. Given this observation, we speculate that the exact number needed may vary by dataset and should be tuned accordingly. Importantly, computation time scales linearly with batch size, and the estimation remains fast, taking only a few seconds, which makes large-scale combinatorial optimization across gene sets feasible. We highlight this capability of the scConcept framework by running a benchmark of a set of publicly available pre-designed gene panels of different sizes (Fig. 5b). For each panel, we estimate the mutual information of the panel with respect to the full transcriptome space for three reference datasets from different tissues, i.e., brain, bone marrow, and ovary. We normalize the MI estimate by the maximum value possible for each dataset, i.e. our estimate of the signal entropy (see Methods). Apart from the obvious effect of the panel size on the MI estimate, we clearly observe (Fig. 5b) that the gene panels designed for brain tissues (*Xenium Brain* and *MERSCOPE Brain 1K*) provide relatively higher information with respect to the panels of the same size for the brain dataset. A similar pattern is also visible for immuno-oncology panels (*CosMX IO 100* and *Xenium IO*), where they appear to be more informative of the whole transcriptome variation of the bone marrow dataset.

**Figure 5:**
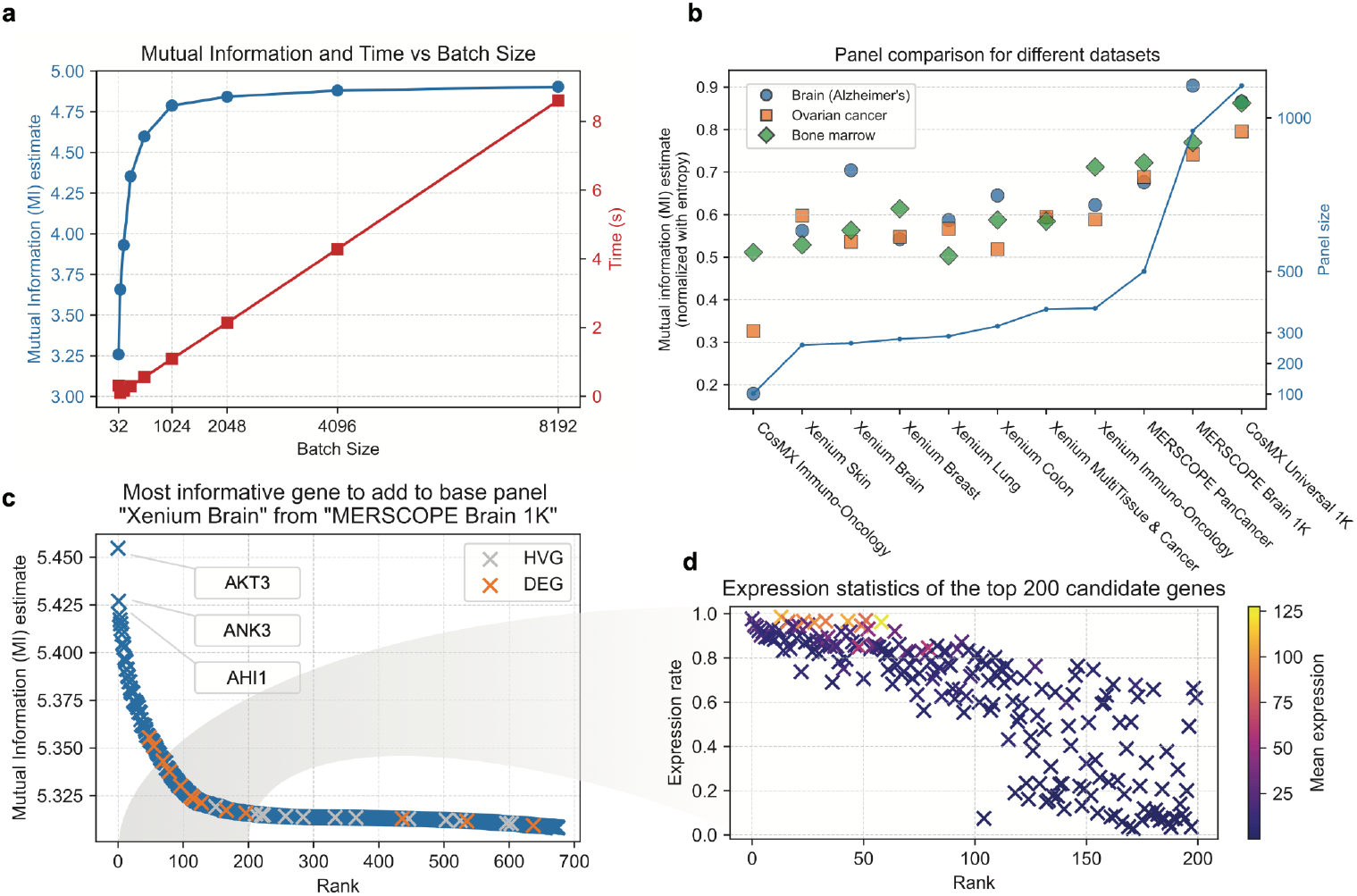
scConcept as a tool for panel design and optimization: (**a**) The accuracy and the computation time of the mutual information estimate between the pair of a Xenium Immuno-oncology panel and the whole transcriptome with respect to the number of cells used for the estimation in a bone marrow scRNA-seq reference dataset. (**b**) The mutual information estimation of different gene panels of different sizes against the whole transcriptome for the three datasets from different tissues and conditions. (**c**) The comparison of the information gain obtained through adding each of the genes from a larger brain panel to a smaller panel in a brain atlas reference dataset. The most informative genes to add are not necessarily among the highly variable or differentially expressed genes. (**d**) The expression statistics (i.e. the portion of cells expressing the gene and the mean expression amount) of the top 200 candidate genes identified in section (c).

We further demonstrate the model’s ability to optimize an existing gene panel by identifying the most informative next gene to add. To this end, we use the snRNA-seq brain atlas from the Alzheimer’s disease dataset and begin with a base panel of 266 genes from the *Xenium Brain* panel. We then consider all genes from the larger *MERSCOPE Brain 1K* panel, which includes 958 genes, and evaluate the mutual information (MI) gain achieved by adding each candidate gene individually to the base panel. The candidate genes are then ranked according to their MI contribution (Fig. 5c). We also highlight the top differentially expressed gene per cell type (resulting in 18 DEGs) as well as the top 20 most highly variable genes from the candidate gene pool. Although DEGs yield higher MI gains than HVGs, they still do not rank among the most informative genes to add to the base panel. This is because being highly differentially expressed does not necessarily provide novel information beyond what is already captured by the existing panel. This is mostly because both DEGs and HVGs are assessed individually for each gene and independently of other genes, which overlooks gene co-expression patterns and mutual dependencies. Examining the list of the top most informative candidate genes identified by the scConcept pipeline reveals several well-characterized functionally active genes in the brain (i.e. *AKT3, ANK3, AHI1*) ^70–76^, many of which display high variability in their expression patterns across the cells and conditions of the data. Looking into the expression statistics of the top 200 candidate genes (Fig. 5d) shows a general correlation between gene expression frequency (i.e., the proportion of cells with non-zero expression), mean expression levels, and their ranking based on the mutual information (MI) gain contributed by each gene. However, despite this correlation, the top-ranked candidate genes are not necessarily those with the highest expression frequency or overall expression levels.

### scConcept maps spatial assays to existing atlases

Single-cell transcriptomics atlases aggregate and integrate data from thousands to millions of individual cells across tissues, organs, developmental stages, and health conditions. These atlases serve as comprehensive reference maps of gene expression, providing spatial, temporal, and functional context for cell types and states. They facilitate cell type annotation in new datasets through label transfer, support the identification of novel or rare cell populations, and enable cross-study and cross-condition comparisons ^77^. One of the main assets of every single-cell atlas is the integrated embedding space, which is optimized to eliminate potential batch effects between different studies while conserving biological relations and similarities. However, most atlases have been built by integrating dissociated whole transcriptome scRNA-seq studies, thus complicating a straightforward integration and usage with spatial datasets that are typically limited to few hundreds of genes. Here we show that scConcept, being a technology-independent cell representor, can be optimized to bridge this gap through learning a technology-independent map to the pre-existing embedding space of current integrated atlases. For this purpose, we consider the Human Lung Cell Atlas (HLCA) ^78^, which combines 49 datasets of the human respiratory system into a single atlas encompassing over 2.4 million cells from 486 individuals. HLCA comes with a publicly available scANVI ^21^ model, which is a conditional Variational Autoencoder (cVAE) based method, that is trained over the top 2000 highly variable genes of all of its collection cells and embeds the cells into a 32-dimensional integrated embedding space.

We propose an atlas mapping adaptation of scConcept (Fig. 6a) where we add a projection layer to translate the pre-trained embedding of the scConcept to the target 32-dimensional space of the HLCA scANVI embeddings. We fine-tune scConcept along with the projection head through the proposed contrastive framework, where we set the second “cell view embeddings” to be the existing atlas embeddings of the corresponding cells at each fine-tuning step (Methods). The combination of gene panel subsetting and the local rank encoding of scConcept results in a technology-independent mapping from the transcriptome space to the integrated latent of the atlas that is unbiased with respect to the input gene panel. We validate the procedure by mapping a healthy Xenium lung sample from a recent spatial transcriptomics study on pulmonary fibrosis ^79^ to the HCLA embedding space. The Xenium panel consists of 343 target genes with manual cell type annotations based on marker gene expression as well as spatial information, available at three different levels of cell type resolutions.

**Figure 6:**
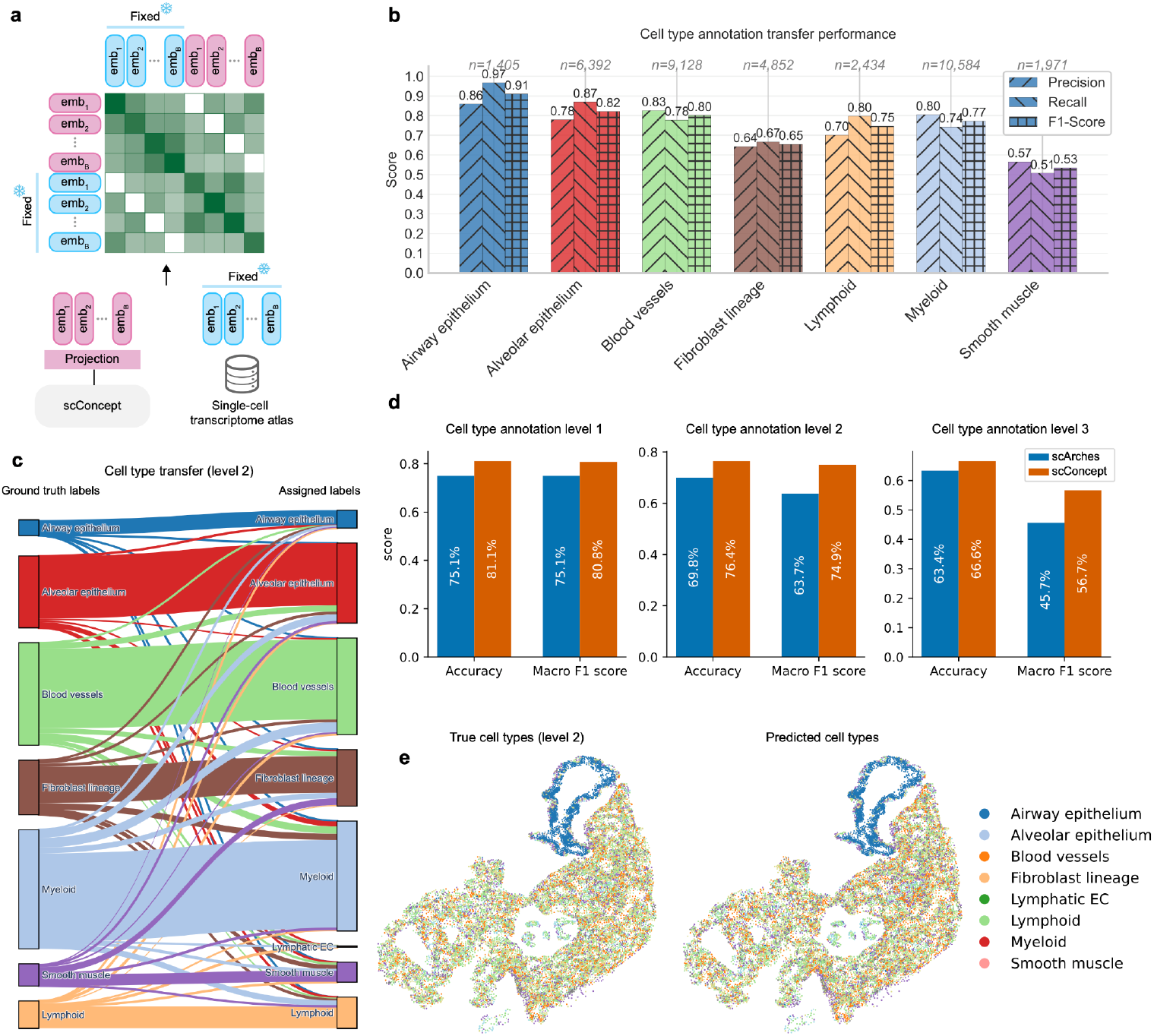
scConcept maps spatial assays to existing atlases. (**a**) The training setting to adapt the scConcept embeddings to the pretrained integrated embeddings of Human Lung Cell Atlas (HLCA). A projection head is added on top of the pretrained base model (scConcept) and then is finetuned over the cells of HLCA with the embeddings from the atlas being fixed. (**b**) The performance of the cell type label transfer through scConcept compared with scArches for the three different hierarchical levels of cell type annotations measured in terms of Accuracy and Macro F1 score. (**c**) The relation between the ground truth cell type labels of the spatial dataset and the assigned labels through projection with scConcept. (**d**) The performance of the cell type label transfer for each cell type in terms of Precision, Recall, and F1-Score. (**e**) The spatial organization of the true and predicted cell types through mapping.

After mapping the spatial dataset of interest to the HLCA reference, we perform cell type label transfer from the HLCA reference to the spatial sample through a KNN classifier with the atlas being the support reference. This allows us to measure the accuracy of the scConcept mapping by comparing the transferred cell type annotations with the labels originating from the manual annotation pipeline. We see a consistent mapping performance across different cell types based on the second annotation resolution (Fig. 6b-c).

While there are numerous methods proposed for mapping dissociated to spatial technologies, none of them learns mappings to the existing embedding space of integrated atlases. The HLCA atlas incorporates scArches ^22^ as a tool to map new unseen dissociated datasets onto the core atlas by training an adaptor added to a reference scANVI model, thereby enabling it to generate a common embedding of the new data. However, as scArches relies on adaptation of the VAE model, any new spatial dataset with a limited gene panel needs to be first zero-padded for the rest of the genes, which can negatively impact the mapping quality. We show that scConcept achieves a substantially higher performance on the cell type label transfer of all three levels of label resolutions, with a larger macro F1 score margin for the most fine-grained level of annotations (Fig. 6d). Examining the spatial organization of true and predicted cell types within a tissue section (Fig. 6e), we find that scConcept’s mapping achieves a biologically coherent recovery of cell type labels, thereby demonstrating its effectiveness in transferring knowledge from existing atlases to new spatial assays.

### scConcept learns robust embeddings across assays and technologies

The challenges of obtaining cellular and subcellular measurements have resulted in a diverse range of technologies and assays with very different protocols and procedures — from cell dissociation to staining procedures and fixation techniques. These differences not only affect the integrity and viability of the cells but also influence the signal characteristics and type of data obtained. The efforts to capture spatially resolved transcriptomes have resulted in another set of new commercialized or academically developed technologies, each offering unique advantages in resolution, throughput, and compatibility with different tissue types. Moreover, targeted spatial transcriptomic technologies require a pre-selection of a subset of genes (i.e., gene panel), which creates an additional layer of variability, as different studies often target different gene panels tailored to specific biological questions or tissue types. This lack of standardization further complicates cross-comparison between assays, making it difficult to benchmark performance or extract new insights across datasets from different platforms. The distinctive training strategy of scConcept combined with the proposed augmentations and input encoding scheme promotes the learning of robust cell representations that are resilient to variations in gene panels and count distributions. As long as the relative expression levels of any subset of genes remain consistent across different technologies, scConcept can successfully map them into a unified embedding space.

We benchmark the robustness of cellular embeddings by evaluating the data integration of different single-cell assays and technologies. In the first case, we measure the integration of single-cell, single-nucleus, and slide-seq assays of the liver samples from metastatic breast cancer biopsies ^80^. Single-nucleus RNA-seq is able to reveal rare cell types and novel cell states, while it does not contain the transcripts in the cytoplasm, which creates a readout shift between the two technologies. Slide-seq, on the other hand, preserves spatial context by capturing transcriptomes directly from intact tissue sections, enabling the mapping of gene expression patterns back to their native histological environment, however it suffers from relatively low capture efficiency and limited sensitivity, which can result in relatively sparse gene expression profiles. In the second case, we analyze the data from samples of ovarian cancer ^81^ obtained from three distinct spatial transcriptomics platforms: CosMx (Nanopore), Xenium (10x Genomics), and MERFISH (Vizgen), encompassing 958, 274, and 140 genes, with 34 genes shared among all three panels. For both cases, we measure the single cell integration benchmark (scIB) ^82^ metrics, including the *Bio Conservation* and *Batch Correction* given the assays as the batch covariates (Methods).

We compare scConcept with the embeddings from scGPT, Nicheformer, and Geneformer as zero-shot methods, while we also compare with PCA as a baseline and scVI ^20^ as a specialized batch correction method (Fig. 7a). scConcept along with the other foundation model approaches can process a variable number of genes as input due to the robust architecture of transformers; however, for the non-transformer baselines, we must either restrict the analysis to the shared genes or apply zero-padding to each dataset to ensure a consistent gene set—a requirement imposed by the computations in both the PCA and VAE frameworks. We tried both approaches and reported the best results (Fig. 7a), which turns out to be subsetting to common genes for PCA and training over the union of genes for scVI by zero padding the unobserved genes in each platform. The results indicate that the zero-shot embeddings of scConcept considerably outperform the corresponding zero-shot foundation models while scConcept+ is able to capture a decent performance compared to PCA and scVI, which is particularly designed as a batch-effect correction method (details of all scores in Supplementary Fig. S8). We further show the UMAP visualizations of the embeddings of the assays for both cases for scConcept+ (Fig. 7b-c) and observe scConcept’s ability to learn technology independent embeddings. It is important to note that while scConcept reduces representation shifts arising from suboptimal training objectives or data modeling, it is not a batch correction method and therefore does not eliminate distribution shifts that reflect genuine differences in cell states or biological variation.

**Figure 7:**
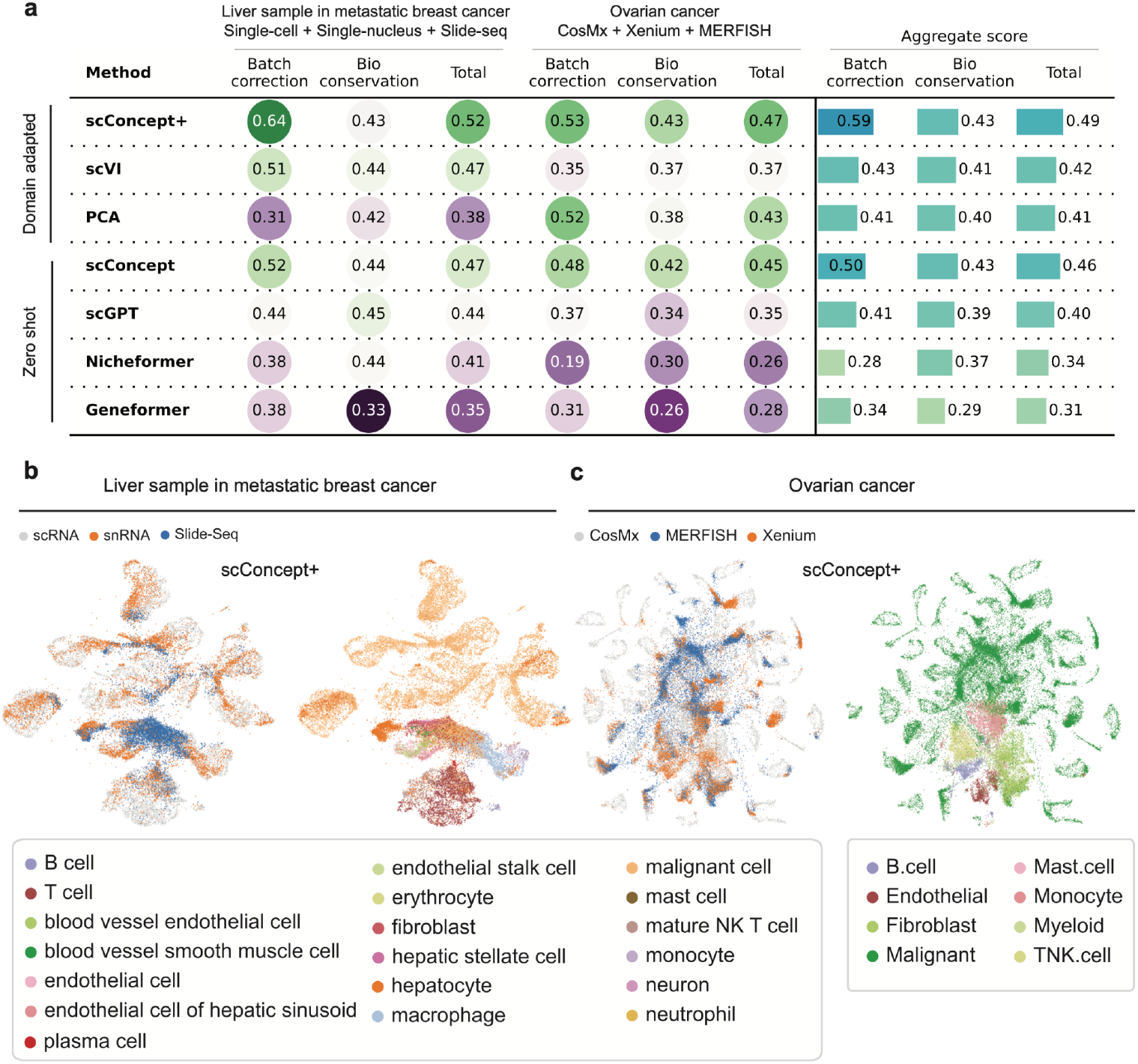
scConcept learns technology-robust cell representations. (**a**) Benchmarking the robustness of cell embeddings through evaluating the data integration of different single-cell assays: 1. Integration of single-cell, single-nucleus, and slide-seq assays of liver sample in metastatic breast cancer. 2. The integration of CosMx, MERFISH, and Xenium spatial assays from human ovarian cancer with different gene panels. We measure the scIB^82^ metrics to evaluate both the biological conservation and the batch effect correction scores given the assay or technology as the batch covariate. Total scores are averages of the two “Batch Correction” and “Bio-Conservation” scores. The scores are aggregated for the two experiments to produce an overall score at the end. (**b-c**) The UMAP visualization of the scConcept+ embeddings colored by assay and cell type labels, showing the robustness of the embeddings with respect to the technology biases.

## Discussion

scConcept represents a significant step toward developing large scale technology-independent single-cell transcriptomic models capable of supporting a broad range of downstream tasks and applications. In contrast to widely adopted masked language modeling approaches, scConcept introduces a novel transformer-based contrastive learning framework, complemented by specialized data encoding and training strategies that are designed to generate robust, technology-agnostic, and information-rich cellular embeddings from transcriptomic data. Our extensive evaluations reveal that scConcept consistently outperforms not only other state-of-the-art foundation models but also specialized, domain-specific methods across a wide array of downstream applications. scConcept demonstrates that rethinking the training paradigm for next-generation single-cell foundation models is crucial for fully leveraging the power of large-scale cellular datasets.

Although multi-omics technologies continue to advance in both accuracy and efficiency, there remains a pressing need for a unified framework to encode cellular measurements across different protocols and platforms. Such standardization would enable the integration and transfer of shared knowledge across diverse assays and technologies. scConcept introduces the first-of-a-kind pretraining framework specifically designed to distill technology-invariant patterns and to generate universal feature representations for the robust characterization and comparison of cell states and identities. Through pre-training on a dissociated corpus and generalizing to unseen sequencing and imaging assays, scConcept provides a potential to bridge experimental heterogeneity, enhance cross-modality interpretability, and facilitate downstream tasks in a consistent and scalable manner.

So far we have trained scConcept on an intentionally limited corpus of only dissociated single-cell transcriptomic datasets, primarily to demonstrate its ability to generalize across platforms and technologies. As a necessary next step, we plan to extend its training to a broader range of experimental modalities to develop a universal cell encoder capable of capturing shared biological patterns across diverse transcriptomic technologies. Currently, scConcept is exclusively focused on transcriptomics. However, applying the contrastive learning paradigm to incorporate additional modalities—such as chromatin accessibility, protein abundance, and spatial organization of cells—could enable even more comprehensive and powerful cellular representations. The underlying principle of training on partial cellular information (“cell views”) in scConcept offers a natural framework for incorporating additional heterogeneous or incomplete data modalities into the training pipeline. Moreover, scConcept introduces a unique cross-scale learning paradigm that directly maps gene-level readouts to cell-level representations. Future work will explore extending this feature to learn higher levels of the biological scale, including tissues and organs.

Despite its strong performance, scConcept has limitations that point toward avenues for future research. First, the performance of the zero-shot model is dependent on the diversity of the pre-training corpus. As seen with a few datasets, tissues underrepresented in the pre-training data benefit substantially from domain adaptation, highlighting the need to expand the pre-training corpus to encompass a wider range of tissues and conditions to improve out-of-the-box generalization. The superior performance of the adapted scConcept+ model across multiple tasks underscores that while powerful, the pre-trained model can suffer from distribution mismatches with new unseen data, requiring an additional adaptation step for optimal results. Moreover, scConcept is an encoder-centric framework, which limits its use in generative applications such as predicting transcriptional responses to perturbations. However, this limitation can be addressed by introducing decoder modules as an auxiliary component on top, without modifying its current training procedure. Such additions could enable the learned representations to be mapped back into the original gene expression space.

In conclusion, scConcept introduces a novel and more effective cell-level contrastive learning objective for pre-training on single-cell transcriptomic data. We demonstrate that scConcept achieves state-of-the-art performance across a wide range of analytical tasks, including cell-type annotation, cross-technology annotation transfer, spatial gene imputation, and data integration. Our results show that scConcept learns genuinely technology-agnostic cell representations, enabling robust analysis across assays with varying count distributions and gene panels. In addition, scConcept unlocks new applications in computational biology, such as a principled framework for gene panel comparison and optimization and a method for mapping new spatial assays onto existing cellular atlases. Overall, our findings establish contrastive pre-training as a powerful and superior strategy for single-cell modeling and foundation model development, laying the groundwork for general-purpose, technology-independent representations of cellular identity.

## Data and Code Availability

All data used are publicly available and are described in detail in the manuscript. scConcept is available as a python package at https://github.com/theislab/scConcept.

## Author contributions

M.B. conceived the project with the help of A.T.L., S.B., and F.J.T.; M.B. implemented scConcept with contributions from A.T.L.; M.B. benchmarked and conducted the analyses with input from S.B., A.T.L., F.S.H.G, and F.J.T.; M.B. wrote the manuscript with contributions and feedback from S.B. and F.J.T.; F.J.T. supervised the project.

## Acknowledgements

We thank Malte D. Luecken and Valentina Beliaeva for their helpful discussion and advice on the atlas mapping use case. Fabiola Curion for helpful comments on the writing. Amir Ali Moinfar for constructive feedback on downstream analyses. Sergei Rybakov, Lukas Heumos, Sunny Sun, and F. Alexander Wolf for their help in setting up and using the LaminDB dataloader. Finally, we thank all Theislab members for their insightful discussions, feedback, and support.

This work was funded by the European Union (ERC, DeepCell - 101054957). It was also supported by the Helmholtz Foundation Model Initiative (HFMI) through VirtualCell. The authors gratefully acknowledge the Gauss Centre for Supercomputing e.V. (www.gauss-centre.eu) for funding this project by providing computing time on the GCS Supercomputer JUWELS at Jülich Supercomputing Centre (JSC). M.B. and S.B. are also supported by the Helmholtz Association under the joint research school “Munich School For Data Science - MUDS”.

## Methods

### Data encoding

The unique and distinct nature of single-cell omics measurements, in particular the scRNA-seq read- outs, requires a proper adaptation to encode such data modality. Raw count modeling ^1^, expression binning ^2,3^, and global gene ranking ^4,5^ are among the most adopted approaches that have been tried at scale for encoding gene expression profiles. Both raw count modeling and expression binning are prone to capturing both technology and dataset-specific batch effects, which limit their generalizability across different assays. Although expression binning tries to improve this by accounting for sequencing depth and library size, it still encodes the unique statistics of the underlying expression distribution (e.g., the shape/skewedness of the count distribution or the proportions of the most abundant count values. Global gene ranking improves this by ordering the genes based on their relative expression across all cells. To achieve that, it calculates mean expressions per gene for each type of assay and increases or decreases the rank of a gene if it is up- or downregulated compared to its average expression in all datasets. However, the averaging stage in this approach is still hugely affected by the batch effect coming from different count distributions across different assays.

Local ranking, on the other hand, is a simple yet solid encoding that orders the genes based on their raw expressions in a single cell; however, it has only been tried in a limited setup through encoding the top 64 expressed genes ^6^. Although this approach may seem a bit more lossy, we show that it is a very robust way of data encoding against any distribution shift across different datasets and also different technologies, including scRNA-seq, snRNA-seq, or various spatial transcriptomic assays. Motivated by its robustness towards shifts across technologies, scConcept employs local ranking for input data encoding, which we have found to be highly effective. For each cell view, genes of nonzero expression are ranked according to their raw expression levels and then combined with positional encodings before being given to the transformer model (Fig. 1d). The local ranking for each cell view provides a robust encoding that mitigates the effects of variations in library size and distribution shifts across different datasets and technologies.

Single-cell RNA sequencing (scRNA-seq) data is typically represented as a cell-by-gene count matrix *X* ∈ ℝ^*N* ×*G*^, where *N* denotes the number of cells and *G* denotes the number of genes. In our setting, we consider a collection of *M* distinct scRNA-seq datasets, each associated with its own count matrix 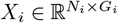, where *N*_*i*_ and *G*_*i*_ correspond to the number of cells and genes in dataset *i*, respectively. Thus, the collection can be expressed as 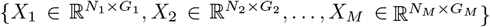.

At each training step, we uniformly sample a mini-batch of *B* = 512 cells of {*cell*_1_, *cell*_2_, …*cell*_*B*_} out of *N* samples for each dataset. Given the count profile of an arbitrary *cell*_*b*_ of the mini-batch, we represent the expression profile of the cell as a sequence of pairs of gene identities and corresponding expression values defined as:

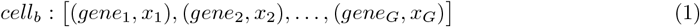

where *x*_*i*_ is the expression amount of *gene*_*i*_ in *cell*_*b*_.

In the next step, as part of the main contrastive approach, we create two positive pairs from a single cellular profile that capture two views of the same cell. To do that, at each training step with a mini-batch of *B* cells, we randomly create two non-overlapping distinct gene panels of *panel*_1_ ={ *gene*_*i*_, *gene*_*j*_,..., *gene*_*k*_ }and *panel*_2_ = {*gene*_*l*_, *gene*_*m*_,..., *gene*_*n*_ }and subset every cell in the minibatch based on both panels, which results in two cell views of the same mini-batch where each captures only a subset of genes for all cells. We have two approaches for creating each gene panel, where we randomly switch between them. The first approach is a pure random panel where we create a panel by randomly selecting a random number of genes from the pool of all genes measured in the dataset from which the mini-batch is drawn. In this case, the panel size is also stochastic, ranging from the full transcriptome to a minimum threshold of 400 genes. The other approach involves selecting from a pre-collected set of pre-designed gene panels from different technology providers. For this approach, we have collected 14 pre-designed gene panels from the official web pages of different technology providers, including: Xenium human brain panel (266 genes), Xenium human breast panel (280 genes), Xenium human colon panel (322 genes), Xenium human immuno-oncology profiling panel (380 genes), Xenium human lung panel (289 genes), Xenium human multi-tissue and cancer panel (377 genes), Xenium human skin panel (260 genes), Xenium prime 5K human pan tissue & pathways panel (5001 genes), MERFISH brain gene panel (140 genes), MERSCOPE pan cancer gene panel (500 genes), MERSCOPE human brain 1K plex (958 genes), CosMx human immuno-oncology panel (100 genes), CosMx universal cell characterization RNA panel (1105 genes), and CosMx Human 6K Discovery Panel (6353 genes).

For each pre-designed gene panel, we also apply a random dropout ranging from zero up to 50% at each training step to bring stochasticity to the fixed pre-designed panels to improve future generalization on unseen panels. Finally, we switch between the two panel creation approaches during training uniformly to cover all possible combinations. Therefore, for each *cell*_*b*_ in the mini-batch, we create two cell views of the following:

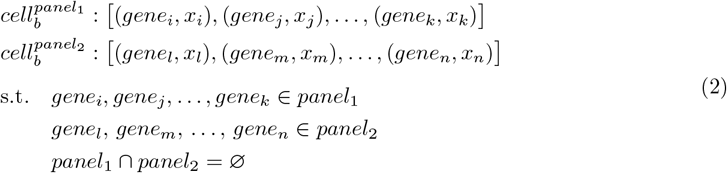

We then process each cell view independently through the Transformer encoder to obtain its corresponding cell view embedding. For this, analogous to natural language processing with Transformers, we tokenize gene identities by assigning a unique token ID to each gene. This is achieved by constructing a one-to-one mapping from gene identities to unique token IDs, thereby representing the *tokenized cell*_*b*_ as:

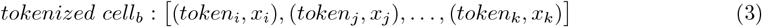

where *token*_*i*_ is the corresponding token id of *gene*_*i*_.

We use rank encoding to encode the expression counts through ordering the gene tokens based on the expression values, and use positional encoding from the original Transformer architecture to encode the sequence order. In particular, we take all the genes with non-zero expression and represent them as:

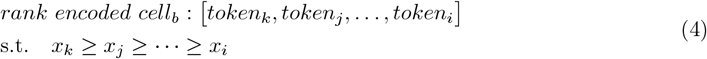

We then use a set of learnable token embeddings of size 512 for each input gene token. The gene embeddings are supposed to capture the identities of each gene and its functional similarity with other genes in the input during training. We set a maximum sequence length of *L* = 1000 gene tokens for cell view input, which results in processing only the maximum top 1000 non-zero highly expressed genes of every input. If the number of expressed genes is less than 1,000, we only process the remaining ones. This constraint comes from the GPU memory available, which creates a trade-off between the mini-batch size, the transformer model size, and the maximum input sequence length. However, a maximum sequence length of *L* = 1000 is a sufficient threshold for most cells sequenced with arbitrary single-cell RNA-seq technologies, given the very sparse nature of the captured transcriptome, which results in very few cells with more than 1000 unique genes being captured.

### Model architecture

We employed a standard Transformer encoder-only architecture composed of 8 layers of multi-head attention blocks. Each block operates with an input and output dimensionality of *dim*_*model*_ = 512 and a feedforward latent dimensionality of *dim*_*feedforward*_ = 1024. Every multi-head attention block features an 8-head self-attention mechanism, followed by a sequence of dropout layers, layer normalizations, fully connected linear transformations, and activation functions. These components are interleaved with residual connections, as detailed below:

We employ a post-normalization architecture, in which layer normalization is applied after the attention and linear layers. A dropout rate of 0.1 is used, and *ReLU* ^7^ serves as the activation function. The above settings result in a Transformer model composed of 16.8 million trainable parameters. For

#### Algorithm 1

Transformer Encoder Block

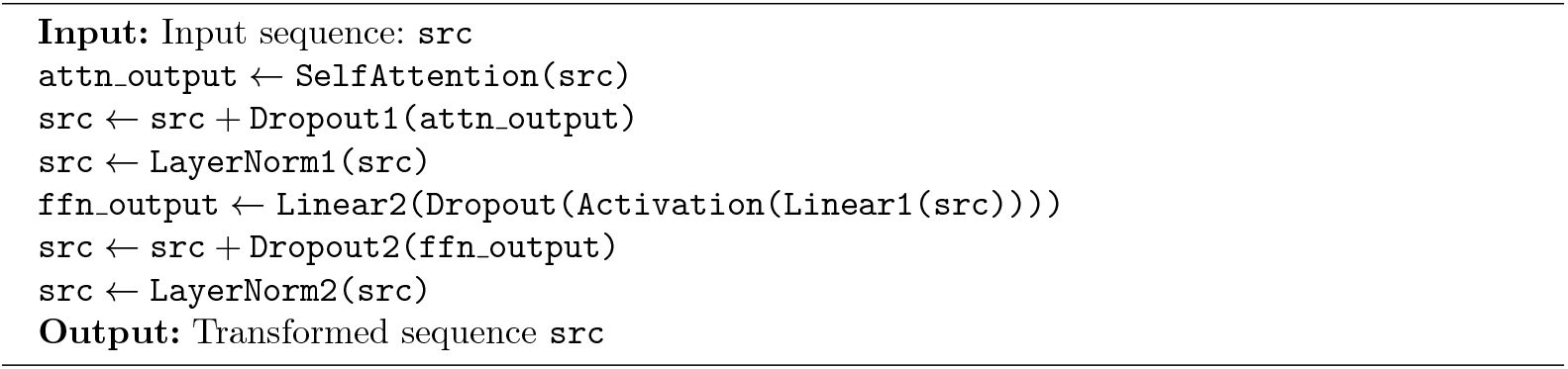

the multi-head self-attention layer, we adopt the Flash Attention implementation ^8^, which substantially reduces memory consumption and accelerates computation by optimizing attention operations through tiling and fused kernels, and enables efficient scaling to longer input sequences.

In order to produce a cell embedding for the whole sequence of gene expression profiles, we define a special class token ([CLS]) ^9^ and add it to the beginning of the input sequence. We therefore take the output embedding of the [CLS] token and define it as the representative cell embedding, which we use to optimize the objective of the contrastive loss. At inference time, we use the output embedding of this token as the output cell representation for all downstream tasks.

### Contrastive learning objective

We use the output of the [CLS] token as the cell representation, which captures the cell’s identity by attending to the entire expression profile sequence. Specifically, we encode the two mini-batch views 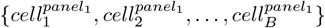 and 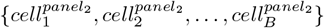, where, for instance, 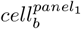 denotes the *b*-th cell in the mini-batch of size *B*, restricted to the gene set defined by *panel*_1_. We obtain the corresponding cell view embeddings of 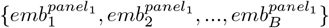 and 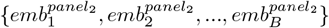 respectively where 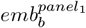 and 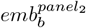 are the embeddings of *cell*_*b*_ restricted to the panels of {*panel*_1_, *panel*_2_ }. In an ideal case, we want the embeddings produced by any subset of genes based on any arbitrary panel to be the same and reflect the constant underlying cell state. This assumption is based on the complex dependency of the whole gene regulatory network, with a high correlation of the co-expression patterns between different sets of genes. Therefore, we expect an ideal network to infer the underlying cell identity through observing any sufficient partial information of the whole transcriptome landscape. To achieve this, we contrast all pairs of cell embeddings across the two views and enforce similar representations for the embeddings of the same cells while maximizing the distance between the embeddings coming from different cells. To achieve this, we calculate the InfoNCE contrastive loss of every positive pair of 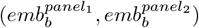:

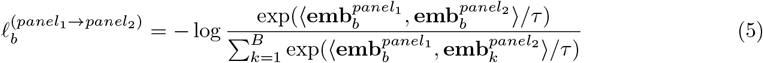

where ⟨., ⟩. denotes the cosine similarity, i.e., ⟨ **u, v** ⟩.= **u**^⊤^ **v***/* ‖**u** ‖ ‖**v** ‖and *τ* ∈ ℝ^+^ represents a temperature parameter that is learned during the training.

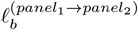 denotes an asymmetric contrastive loss by contrasting each cell from *panel*_1_ to every cell from *panel*_2_ ((*panel*_1_ → *panel*_2_)). Similarly, we construct the complementary asymmetric loss term for *panel*_2_ *panel*_1_. The final objective is defined as the symmetric average of these two loss terms, optimized over all positive pairs within each mini-batch.

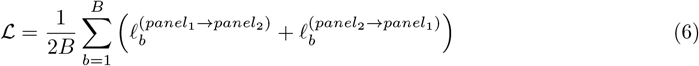

The above is a symmetric InfoNCE contrastive loss that contrasts each cell of every panel with all the cells of the other panel.

As we represent each cell based on the non-zero expressed genes, the absence of a gene in a cell view could either be caused by not being included in the selected panel or by the zero expression of the gene in that specific cell. We observe that this dual meaning of the absence of a gene can encourage the model to be sensitive to the presence or absence of a gene in the input, and therefore result in similar embeddings for different cells with different expression profiles, but only with the same gene panel. Therefore, we introduce a modified version of the contrastive loss in Eq. 5 where we contrast a cell view embedding not only with the embeddings of the opposite view but also with the embeddings from the same view (same panel):

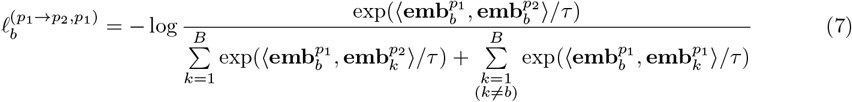

where *p*_1_ and *p*_2_ represent *panel*_1_ and *panel*_2_ respectively. The first term in the denominator contrasts a cell with negative pairs of the different view, while the second term contrasts it with the other cells of the same view and panel. We finally use the corresponding symmetric version of the modified loss for the final training:

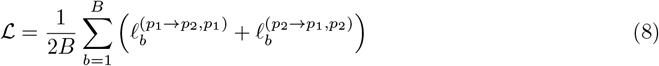

Through this modification, we make sure that what the model learns is based on the order of the gene expressions and not the presence or absence of a gene in the panel. For the stability of the training, we use the easier cross-view contrastive loss (Eq. 5) for the warm-up phase of the training (the first 20,000 training steps) and then permanently switch to the second, more challenging objective (Eq. 7) for the rest of the training, which shows a consistent convergence behavior.

### Training setting

We trained our model using the Adam optimizer with a learning rate of 10^*-*4^. A linear warm-up strategy was applied for the first 20*K* steps to stabilize optimization, in which the learning rate is gradually increased from zero in the beginning to 10^*-*4^ at step 20*K*, after which the learning rate remains constant. To facilitate stable learning, we use the cross-view loss (Eq. 5) for the warm-up phase and then switch to the main loss (Eq. 7) afterwards. Training is performed in a multi-GPU setting, where batches are distributed across available GPUs through the Distributed Data Parallel (DDP) strategy to improve throughput, which allows for larger mini-batches, context windows, and model size. However, the model itself is not distributed across devices as it fits on every single GPU. This setup ensured efficient utilization of hardware resources while maintaining consistency in gradient updates. We train the model on 4 x H100 Nvidia GPUs for 48 hours, which is equivalent to over 1 million gradient updates.

### Data loading and sampling strategy

The large pre-training data corpus, such as the one used in our case, does not fit in a compute node’s memory and therefore needs to be loaded from disk during training. At each step of training, we sample a random mini-batch of cells each time from a single dataset as part of our sampling strategy. A contrastive approach relies on the diversity of the cells contrasted to each other at each training step. Therefore, having true, real-time random access to all cells from a pool of a large number of datasets is essential for training. The data loading tools that only enable pseudo-shuffling (i.e., splitting the data into chunks that fit into memory and pre-shuffling the chunks once) do not satisfy our criteria. We use the LaminDB (https://lamin.ai/) framework, which offers efficient data loading and random access to all cells of the pre-training data through working with native h5 anndata objects (i.e. h5ad files).

### Inference time domain adaptation: scConcept+

Despite the large pre-training data, the diversity and heterogeneity of the transcriptional landscape across donors, conditions, and disease can go beyond the pre-training corpus, which may result in out-of-distribution scenarios for new datasets. Here, we propose the inference-time model adaptation approach, which adapts the pre-trained scConcept model for unseen datasets at test time. To achieve this, for any new given target dataset, we load the pre-trained model scConcept and continue training it with the main contrastive loss (Eq. 7) over the random mini-batches sampled from the target unseen dataset. This results in a self-supervised adaptation of the base model to which we refer as scConcept+. This type of model adaptation is different from task-dependent fine-tuning of models, which is done for a specific task, such as cell-type annotation, where the base pre-trained model is fine-tuned to solve a target task of interest by optimizing over supervised labels. The number of adaptation steps relies on the target dataset size; however, for the datasets of fewer than 100K cells, we typically find that an adaptation of 50,000 steps (corresponding to 50,000 weight gradient updates) is sufficient and optimal for capturing the heterogeneity of the unseen target data (Supplementary Fig. S1). Unlike the pre-training phase that is performed over multiple GPUs with large mini-batch size of *B* = 512, we do a lightweight adaptation using a mini-batch size of *B* = 128 which fits on a single GPU memory and is generally quick compared to the pre-training phase (e.g. takes 54 minutes for a target dataset of about 43*K* cells for a total of 30*K* adaptation steps on a single H100 Nvidia GPU.

### scConcept for gene panel mutual information estimation

Mutual information (MI) is an information-theoretic measure that quantifies the reduction in uncertainty about one random variable gained from knowledge of another. In other words, it captures the amount of information shared between two random variables and is formally defined as:

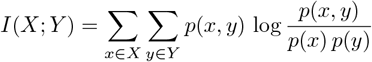

where *X* and *Y* are arbitrary random variables, and the degree of dependence between them determines the magnitude of their mutual information. This definition is not restricted to univariate random variables but naturally extends to high-dimensional multivariate variables, allowing the measurement of mutual information between sets of univariate random variables. Given a probability distribution over the gene expression profile *G* = {*G*_1_, *G*_2_,..., *G*_*n*_}, we can define the mutual information between any two subsets of genes, *X*_1_ ⊆ *G* and *X*_2_ ⊆ *G*. Owing to the strong interdependencies among genes in the transcriptome, driven by complex regulatory networks, mutual information provides a principled way to quantify how much information a gene panel *X*_1_ conveys about another gene panel *X*_2_ under a given gene expression distribution. Calculating the exact mutual information between two gene panels requires knowledge of both their marginal and joint distributions, which are generally inaccessible for real-world data.

Contrastive learning provides a well-established framework for estimating mutual information between sets of high-dimensional random variables at scale ^10^. Building on the formulation in ^10^, we demonstrate that scConcept can be applied to estimate the mutual information between arbitrary subsets of gene panels. Specifically, as discussed earlier, scConcept optimizes the InfoNCE contrastive loss over pairs of gene panels *X*_1_ and *X*_2_:

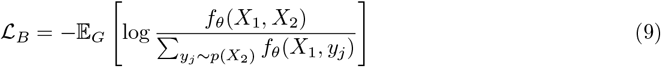

where *f*_*θ*_(.,.) is the similarity score between the extracted embeddings of scConcept. As demonstrated in ^10^, the optimal similarity function that minimizes the InfoNCE contrastive loss is proportional to the ratio between the joint distribution of the gene panels and the product of their marginal distributions, given by:

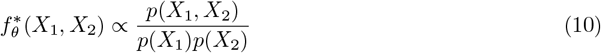

As shown in ^10^, substituting the optimal similarity function 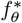 into Eq. 9 yields a lower-bound estimate of the mutual information between the two gene panels:

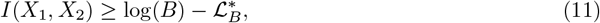

where *B* denotes the mini-batch size used to approximate the expectation in Eq. 9.

Although optimizing scConcept with the InfoNCE contrastive objective (9) via stochastic gradient descent may not reach the global minimum of Eq. 9, it still produces a valid, albeit looser, estimate of the mutual information, since Eq. 11 also holds for suboptimal similarity functions *f*_*θ*_ that yield higher values of ℒ_*B*_.

scConcept, when pretrained on a large single-cell transcriptome database, already achieves a low contrastive loss on previously unseen datasets, thereby providing a lower-bound estimate of the mutual information between any two subsets of genes (gene panels) as in Eq. 11. However, further adaptation of the model to the target dataset by scConcept+ via continuing to optimize the contrastive loss can reduce the loss even further, yielding a tighter and more accurate estimate of the mutual information.

As shown in ^10^, the lower-bound estimate in Eq. 11 is theoretically independent of the batch size (*B*), but its accuracy improves as *B* increases. Therefore, it is advisable to determine the minimum required batch size that still provides a reliable estimate of the mutual information for any given target dataset.

### Pretraining data

scConcept is pre-trained on over 33 million gene expression profiles from 241 datasets, encompassing 13,035 donors across various tissues and organs, sourced from the CellxGene census collection32. To evaluate the improvements brought by the proposed pre-training approach, we deliberately restricted our pre-training corpus to the same CellxGene release version used by scGPT^3^. This results in around 33 million cell profiles, which is comparable to the Geneformer^4^ training data in terms of the number of cells. By selecting a pre-training corpus that exclusively includes whole transcriptome sequencing technologies, we also demonstrate that the proposed approach can seamlessly generalize to other technologies, such as image-based spatial transcriptomics, that were not part of the training data.

We acquired the set of all Homo Sapiens datasets encompassing 36, 227, 903 unique cells in total from the CellxGene portal as of release 15 Dec 2023 using the CellxGene census API available at https://chanzuckerberg.github.io/cellxgene-census/index.html, where we downloaded all the datasets in their native Anndata objects. We then filtered for only the most abundant 7 assays of “10x 5’ v2”, “10x 3’ v3”, “10x 3’ v2”, “10x 5’ v1”, “10x 3’ v1”, “10x 3’ transcription profiling”, and “10x 5’ transcription profiling”, which resulted in 272 unique datasets. We then split the data into training and validation sets of 245 and 27 datasets with sizes of 33, 436, 305 and 1, 287, 784 cells, respectively. Although all selected assays are based on whole-transcriptome technologies, the number of genes available in each dataset varies considerably. This variation arises from differences in sequencing depth, alignment methods and software versions, gene-mapping strategies, and optional filtering steps that restrict analyses to protein-coding transcripts. As a result, the reported gene counts range from 15,991 to 60,664 across all the datasets. We only filter for the 19,331 protein-coding genes and drop the rest of the non-coding genes for all datasets. We observed that training over all the reported gene set (protein and non-protein coding transcripts) did not result in any significant improvement in the cell-type annotation performance; however, it significantly increases the convergence time of the model due to less frequent updates to each gene embedding given the fixed context window size.

### Datasets for downstream analysis

For all downstream tasks, we use datasets that were not included in the pre-training phase, thereby simulating a real-world scenario where the model is applied to previously unseen data during inference. To enable efficient evaluation across multiple models and datasets, we adopted a subsampling strategy: for any dataset containing more than 100,000 cells, a random subset of 20,000 cells was selected for evaluation. The selection procedure is entirely random and does not introduce any bias with respect to any class labels or covariates in the data. In all datasets, we retain the complete set of genes, irrespective of their inclusion in the gene vocabulary of scConcept. This choice allows fair comparison with other foundation model approaches that accommodate larger gene vocabularies. The full gene set is likewise used for all baseline methods, including normalized counts, PCA, CellTypist, Tangram, scArches, and scVI.

#### Bone Marrow

This dataset was released as part of the NeurIPS 2021 Multimodal Single-Cell Data Integration open problems challenge (https://openproblems.bio/events/2021-09_neurips) ^11^. The version used in our experiments comprises 42,492 mononuclear cells derived from 12 healthy human donors, collected across three experimental sites. Although the data originate from a multiome assay—jointly profiling gene expression (GEX) and genome-wide chromatin accessibility (ATAC)—only the gene expression modality was used for annotation. The dataset encompasses 22 distinct cell types and includes expression measurements for 13,431 genes. For the annotation task, data from the first site were designated as the reference set, while data from the third site were used as the query set.

#### Brain (Multiple Sclerosis)

This MS dataset is available at EMBL-EBI https://www.ebi.ac.uk/gxa/sc/experiments/E-HCAD-35, which scGPT^3^ also adopted as part of its cell-type annotation evaluation. This dataset contains nine healthy control samples and twelve MS samples. The control samples were used as the reference set, while the MS samples were held out as the query set for evaluation. We use the same pre-processed data as in scGPT, which excludes three cell types—B cells, T cells, and oligodendrocyte B cells—that were present only in the query dataset. After filtering, the reference set contained 7,844 cells and the query set contained 13,468 cells. Furthermore, highly variable genes (HVGs) were selected to retain 3,000 genes. Cell type annotations from the original publication are used as ground truth for evaluation.

#### Skeletal Muscle Aging Atlas

This dataset is a human skeletal muscle aging atlas, created by profiling the transcriptomes of 90,902 single cells and 92,259 single nuclei from the intercostal muscles of 15 human donors spanning the adult human lifespan ^12^ and is accessible from the CellxGene data portal. These donors were divided into two groups: eight young individuals (approximately 20–40 years old) and seven aged individuals (approximately 60–75 years old). Eight random donors were selected as the reference set, and the remaining seven donors were used as the query set. The combined use of single-cell and single-nucleus RNA sequencing allowed researchers to identify and annotate 40 major cell populations within the muscle tissue.

#### Seattle Alzheimer’s Disease Brain Cell Atlas

The Allen Brain Map atlas (https://portal.brain-map.org/) provides paired datasets from the middle temporal gyrus of aged human donors, including single-nucleus RNA sequencing (snRNA-seq) from 84 individuals and multiplexed error-robust fluorescence *in situ* hybridization (MERFISH) from a subset of 27. The snRNA-seq dataset profiles 36,412 genes, whereas the MERFISH panel targets 139 genes. All data were downloaded from https://registry.opendata.aws/allen-sea-ad-atlas.

#### Ovarian Cancer Assays

A 10x scRNA-seq whole-transcriptome sample (18,082 genes) paired with two 10x Xenium spatial assays from an ovarian cancer sample were obtained from the publicly available 10x data portal (https://www.10xgenomics.com/datasets/). The two spatial assays use the 5K human pan-tissue panel (5001 genes) and the immuno-oncology profiling panel (380 genes), respectively. All datasets originated from a single donor and comprise 16 harmonized cell types.

#### Human Lung Cell Atlas (HLCA)

This is a comprehensive atlas integrating 49 datasets with over 2.4 million cells from 486 individuals ^13^. The foundation of the atlas is the “HLCA core” a detailed healthy reference built by integrating 14 datasets containing 584,444 cells from 107 individuals with diverse demographic backgrounds. The core donors exhibit a wide range of characteristics, including age (10–76 years), sex (60% male and 40% female), smoking status (52% never-smokers), and harmonized ethnicity (65% European). The atlas is from 166 tissue samples using a variety of experimental protocols and sequencing platforms to capture cellular landscapes across different anatomical locations of the lung. The HLCA study also provides an already integrated scANVI model that projects cells into a 32-dimensional embedding space based on the top 2000 highly variable genes.

#### Xenium Lung Sample

Xenium Image-based spatial transcriptomics dataset comprising 45 lung tissue samples from 35 unique individuals, including nine unaffected donors and 26 participants with pulmonary fibrosis (PF). The assay is performed on a Xenium panel with 343 target genes. The data can be accessed from the Gene Expression Omnibus (GEO) database with the accession number GSE250346. The data from a single healthy donor (accession number GSM7990532) have been selected for the application of mapping to the HLCA atlas.

#### Multi-assay Liver Samples of Metastatic Breast Cancer

The original study of this data ^14^ generated a multi-modal map of metastatic breast cancer by profiling 67 tumor biopsies from 60 patients, with 30 processed by scRNA-seq and 37 by snRNA-seq. A comparison of these methods showed that snRNA-seq yielded a higher number of observations and features per observation and more faithfully captured malignant and stromal cells, whereas scRNA-seq captured a higher fraction of immune cells. For 15 of these biopsies, matching spatial data was generated using Slide-seq, a technology that profiles the whole transcriptome using 10-*µ*m beads. Slide-seq provided a similar number of observations (beads) per sample as the single-cell methods, but detected far fewer features per observation. For our evaluations, we have only subsetted the data to include the biopsies from the liver tissue for all three assays. The data for all the technologies are available at the CellxGene portal.

#### Ovarian Cancer Multi-platform Spatial Data

This spatial dataset was collected using three different platforms, with the CosMx Spatial Molecular Imager (SMI) being the primary technology used to generate the main data, measuring 958 genes across 491,792 cells from 94 tumors. For technical comparison and validation, the original study also used in situ sequencing (ISS) via the Xenium platform to profile 274 genes in 32 tissue cores and multiplexed error-robust fluorescence in situ hybridization (MERFISH) to profile 140 genes in four fresh-frozen tissue sections. The data is also publicly available at Single Cell Portal (SCP2640, SCP2641, and SCP2650).

### Details on downstream tasks and evaluations

#### Cell type annotation

To annotate cell types in a query dataset using a reference dataset, we first embed both datasets into a shared latent space using scConcept (or any competing method). For each cell in the query set, we then identify its *k* = 10 nearest neighbors among the embedded reference cells using Euclidean distance, and assign the most frequent cell type among these neighbors as the predicted label. For this section, we intentionally choose to use a KNN classifier to directly evaluate the structure of the embedding space and avoid the effect of any hyperparameter tuning of a linear or multi-layer neural network. For the PCA baseline, we first normalize the total counts of each cell to 10,000, followed by a log1p transformation (*log*(*x* + 1)). We then evaluated embeddings constructed from the top 32, 128, and 512 principal components, and selected the best-performing configuration. This corresponded to 128 PCs for the bone marrow and skeletal muscle datasets, and 32 PCs for the brain MS dataset (Supplementary Fig. S2). However, due to the substantial variation in performance across different choices of principal components, selecting an appropriate number of PCs is non-trivial in practice—especially when annotating an unlabeled target dataset where groundtruth labels are unavailable for hyperparameter tuning. This limitation highlights a drawback of relying on PCA embeddings for downstream tasks. In contrast, CellTypist is a supervised linear neural network explicitly trained for cell-type prediction. Therefore, for this baseline, we did not apply a *k*NN classifier on any embedding space; instead, we directly trained the provided model and inferred predictions using the official Python API: https://github.com/Teichlab/celltypist.

We evaluate cell type prediction performance using both accuracy and macro F1 score. To compute the macro F1 score, we first calculate precision, recall, and F1 for each cell type present in the query set (cell types appearing only in the reference set are ignored). We then take the unweighted mean of these per-class F1 scores, i.e., the macro-average, which assigns equal importance to all cell types regardless of their frequency. Consequently, a higher macro F1 score reflects consistent performance across both abundant and rare cell types, rather than being biased toward majority classes.

#### Dissociated to spatial cell type transfer

scGPT, Geneformer, and Nicheformer are transformer-based models that inherently support a variable number of genes in the input without requiring any adaptation for unmeasured genes. For the Normalized count and PCA baselines, we either have to fill in the unmeasured genes of the Xenium panel with zeros for the query set or only operate over the shared set of genes between the reference and query sets. The former approach resulted in a very poor performance, which is expected as the zero inflation of tens of thousands of genes distorts the similarity distances both in the original count space and the PCA space. Therefore, we only evaluate with the second approach and report the performance of cell-type transfer by operating only over the shared genes.

#### Spatial gene imputation

For evaluating the imputation of the pseudo-spatial dataset, we iterate over the following publicly available panels from the different technology providers and subset the query set based on each panel, and impute the rest of the genes not included in the panel:

1. Xenium human immuno-oncology profiling panel (380 genes)
2. Xenium prime 5K human pan tissue & pathways panel (5001 genes)
3. MERSCOPE pan cancer gene panel (500 genes)
4. CosMx human immuno-oncology panel (100 genes)
5. CosMx universal cell characterization RNA panel (1105 genes)

For the real pairs of spatial and dissociated references, we can only evaluate the imputation by splitting the genes of the query set into train and test sets and evaluating the imputation over the test split. We do a 6-fold evaluation based on different sets of held-out genes for each dataset.

As many genes may not have any variation across a dataset, for sparse datasets, we only evaluate over the set of highly variable genes (HVGs) of the held-out genes. Therefore, we use the top 1,000 HVGs for the pseudo spatial experiment, 100 top HVGs for the 5k prime Xenium, and 50 top HVGs for the immuno-oncology Xenium ovarian cancer samples.

For comparison with normalized count space, we take the shared genes of the query set and the scRNA-seq reference and normalize both by first normalizing each cell by total counts over all genes, so that every cell has the same total count of 10,000 after normalization. We then perform log1p transformation on both sets. Then, for each cell in the query set, we find the top (K=10) closest neighboring cells in the reference set and perform k-nearest neighbors regression to impute all the genes not in the spatial panel but available in the whole transcriptome reference. For the PCA we run the same pre-processing except that we take the top 30 principal components (PCs) after normalization and find the closest neighbors in the reduced PCA space.

For evaluating the imputation performance, we compute the Pearson correlation coefficient (PCC) and the Spearman’s rank correlation coefficient score between the predicted expression and the totalcount normalized and log1p-transformed ground truth expression for each gene across all cells, and then aggregate the scores across the held-out genes. We calculate the PCC score for every gene *g* as:

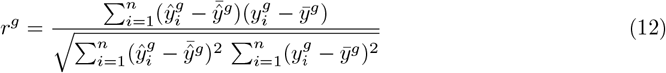

where 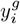 and 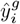 are the ground truth expression (normalized and log1p transformed) and the imputation predictions of gene *g* in cell *i*. Next, we calculate the overall PCC across all held-out genes as 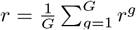.

Since the count distributions of the reference and query sets may not be aligned, we additionally assess imputation performance using Spearman’s rank correlation coefficient. This metric computes the PCC on the ranked values of the predicted and ground-truth expression levels, and the resulting values are then averaged across all held-out genes. While Pearson’s correlation assesses linear relationships, Spearman’s rank correlation assesses monotonic relationships (whether linear or not).

#### Mapping to a pre-existing atlas

To align new spatial datasets with the embedding space of a pre-trained atlas, we fine-tune scConcept jointly with an additional projection head. The goal is to learn a mapping from the high-dimensional scConcept embedding space (512 dimensions) to the low-dimensional integrated embedding space of the HLCA atlas (32 dimensions). During fine-tuning, at each iteration, we randomly sample a mini-batch of atlas cells and construct a single view by subsetting their genes to an augmented panel (using the same panel generation procedure as in pretraining). We then rank-encode the view and compute scConcept embeddings (512 dimensions) for this view and pass them through the projection head to obtain 32-dimensional embeddings aligned with the atlas latent space. Next, we construct a similarity matrix between the projected embeddings and the corresponding pre-trained atlas embeddings of the same cells, and optimize a contrastive loss based on these similarities. During backpropagation, the atlas embeddings remain fixed, while the weights of both scConcept and the projection head are updated to learn the mapping from any gene panel to the target atlas embedding manifold.

#### Technology integration

We use the single-cell integration benchmarking (scIB) pipeline ^15^ to quantify the capability of a cell embedding in correcting for the technology-induced biases and batch effects while conserving the biological variation. We use the official implementation of the scIB pipeline available at (https://scib-metrics.readthedocs.io/). Through this benchmark, we measure the set of biological conservation scores, including Isolated labels, K-Means ARI, K-Means NMI, and Silhouette Label, along with the set of batch effect correction scores, including Batch removal adapted silhouette (BRAS), K-nearest-neighbor batch-effect test (KBET), and the Graph Connectivity score. The details of each of the metrics are available in ^15^.

## Supplementary materials

**Fig. S1:**
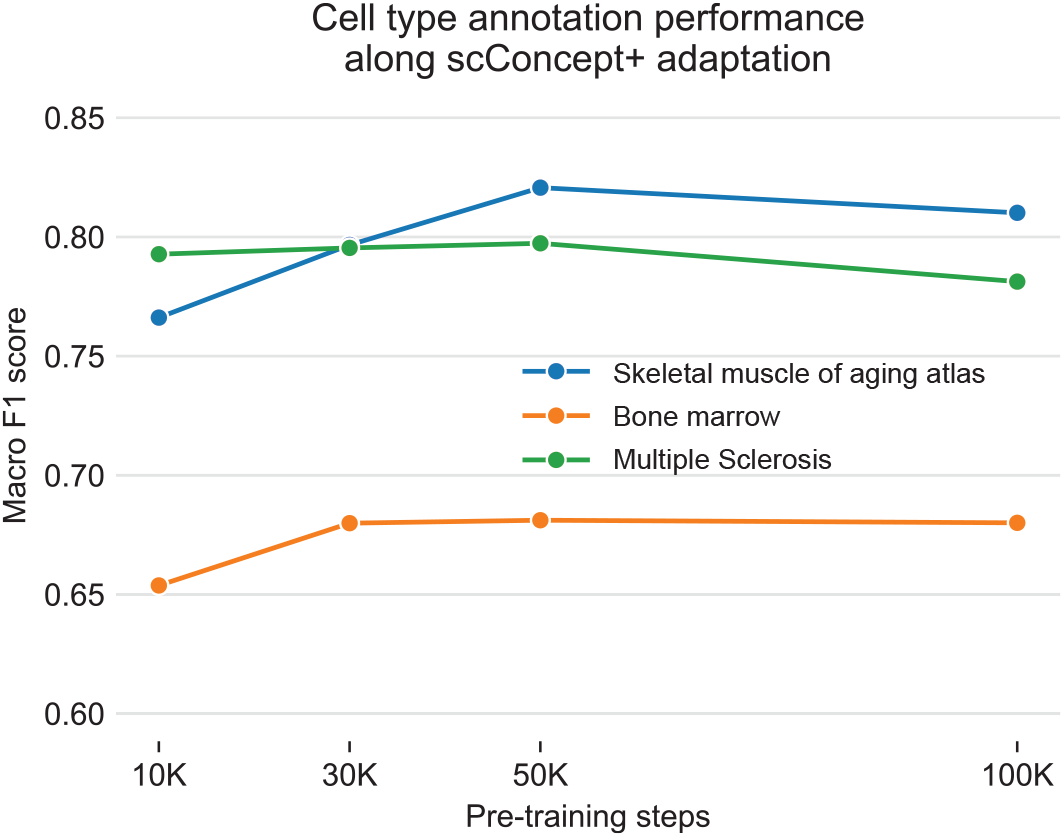
The cell-type annotation performance along the adaptation of scConcept+ with respect to the number of training steps of optimizing the self-supervised contrastive objective over the three target datasets separately. This analysis shows that 50,000 adaptation steps seem to be optimal across different datasets of size less than 100,000 cells. To have an estimate: for a dataset of size 100,000 cells, each cell will be observed *B* = 128 times for an adaptation of 100,000 steps with a mini-batch size of *B* = 128.

**Fig. S2:**
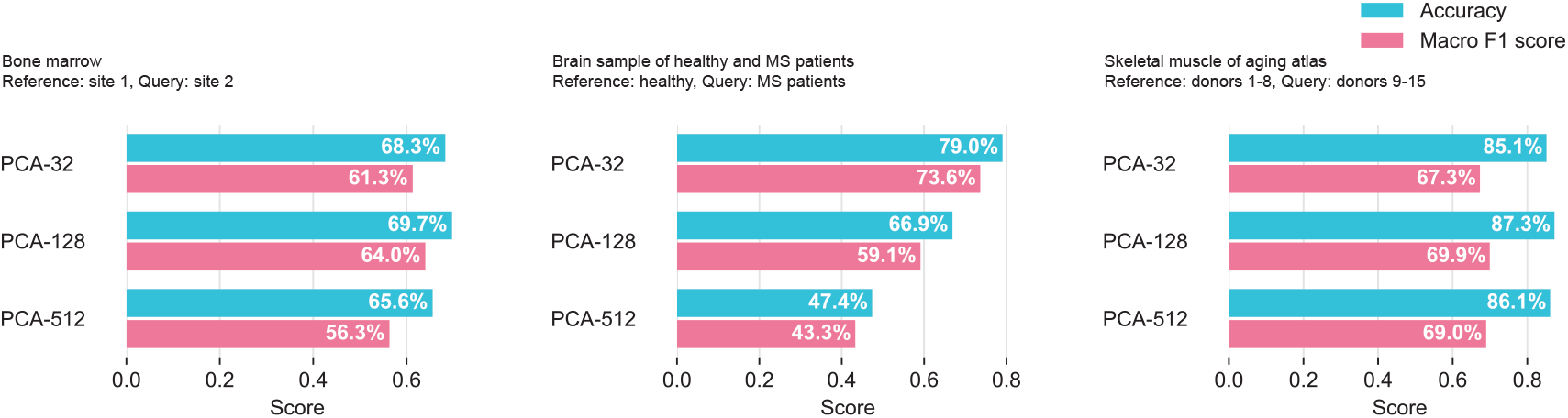
The performance of the cell type annotation using different numbers of principal components of the PCA. Accordingly, 128 PCs perform the best for two of the datasets (bone marrow dataset and Skeletal muscle dataset), and 32 PCs perform the best for the brain sample MS dataset. We picked the best performing one for comparison with scConcept.

**Fig. S3:**
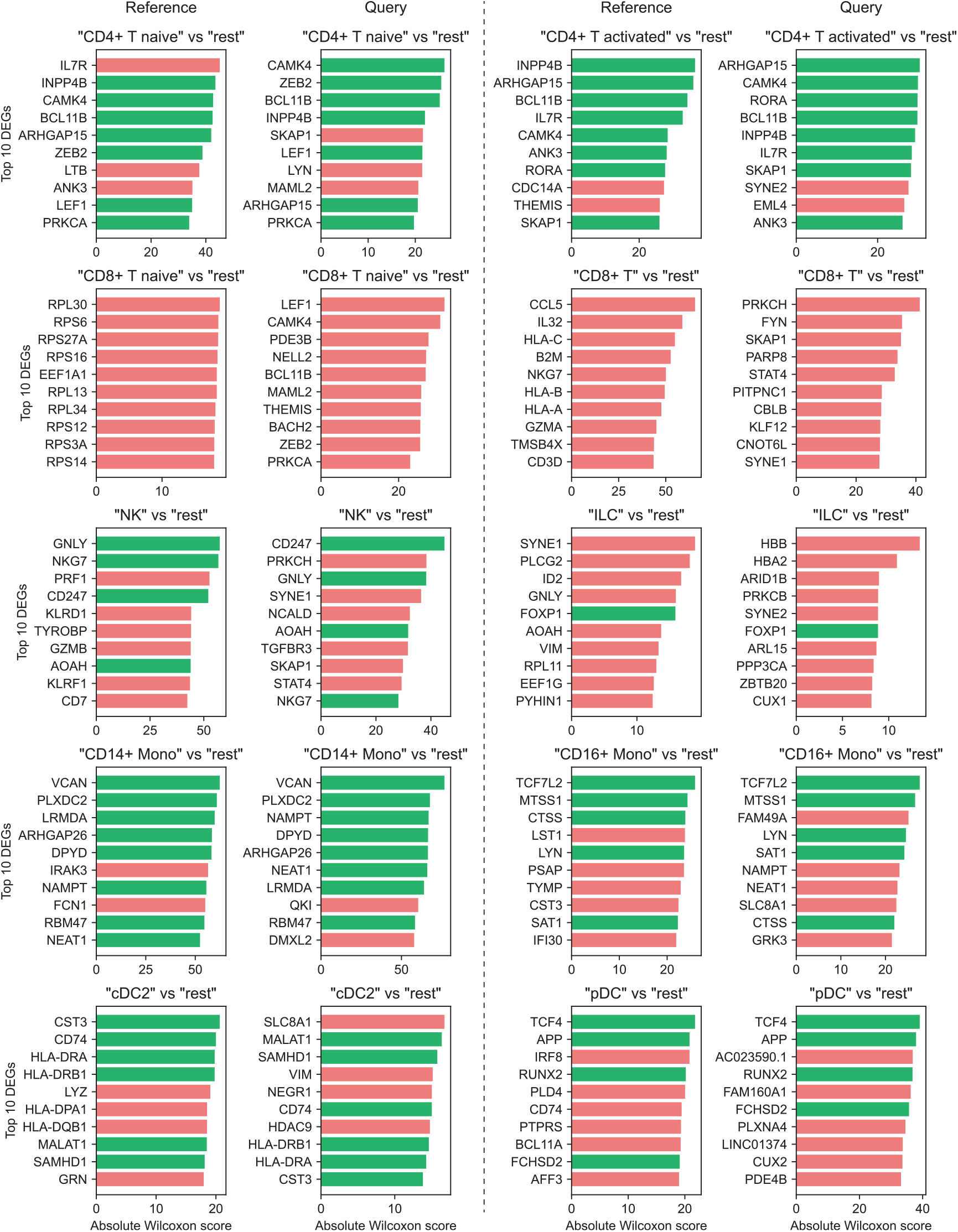
The top differentially expressed genes of each cell type group were calculated in the reference and query sets separately for the bone marrow dataset, sorted by the absolute Wilcoxon score. The genes that are among the top differentially expressed genes of both the reference and query sets are colored in green; otherwise colored in red. (Continued on next page)

**Fig. S4:**
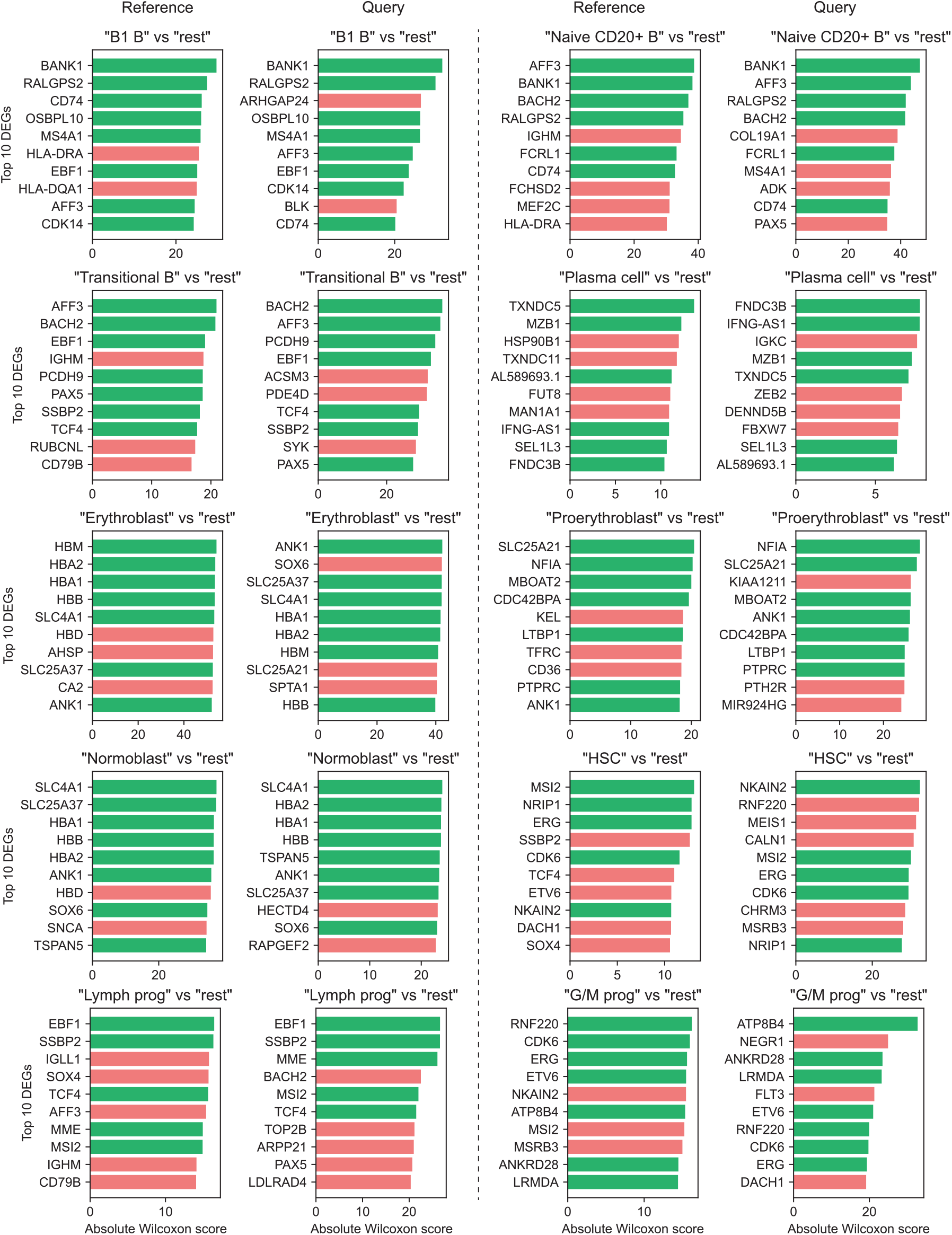
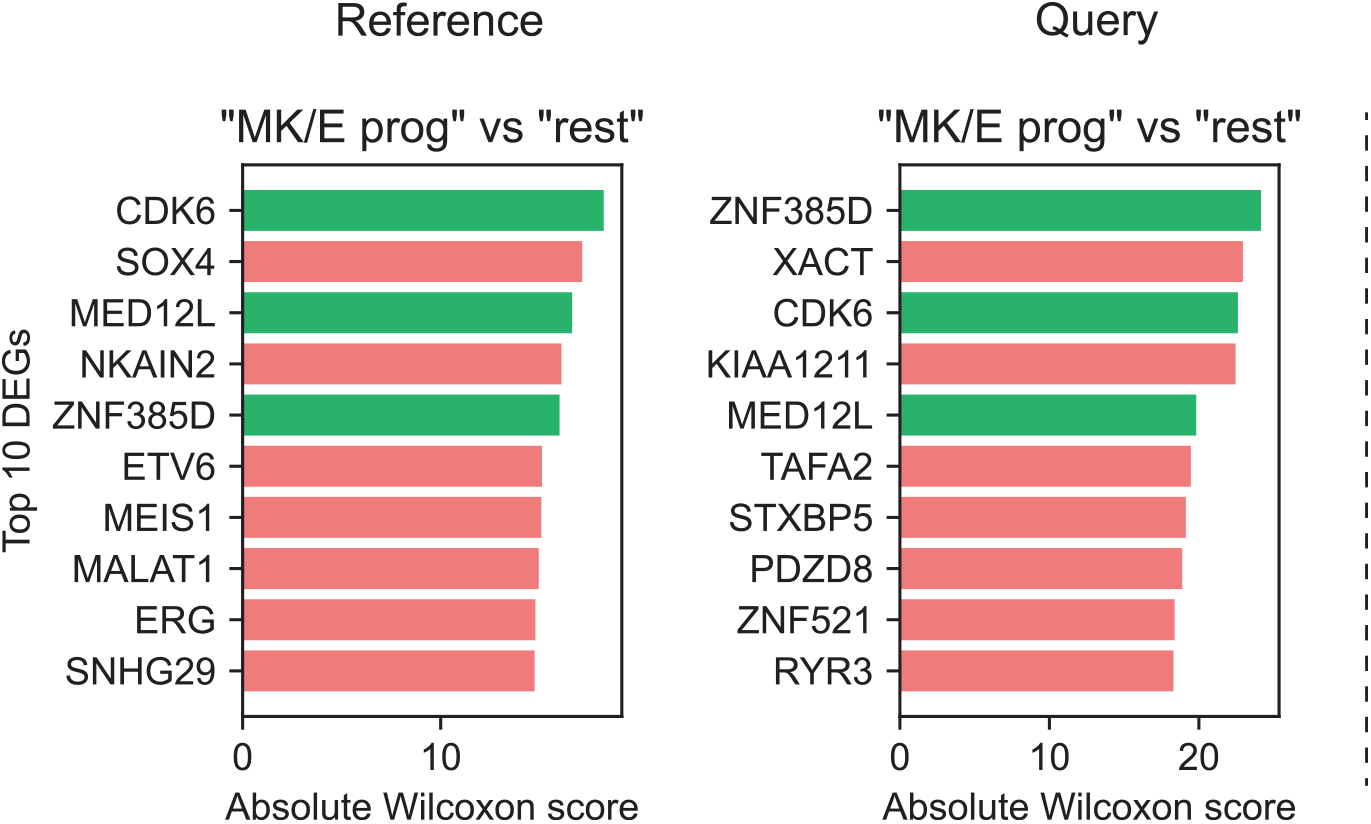
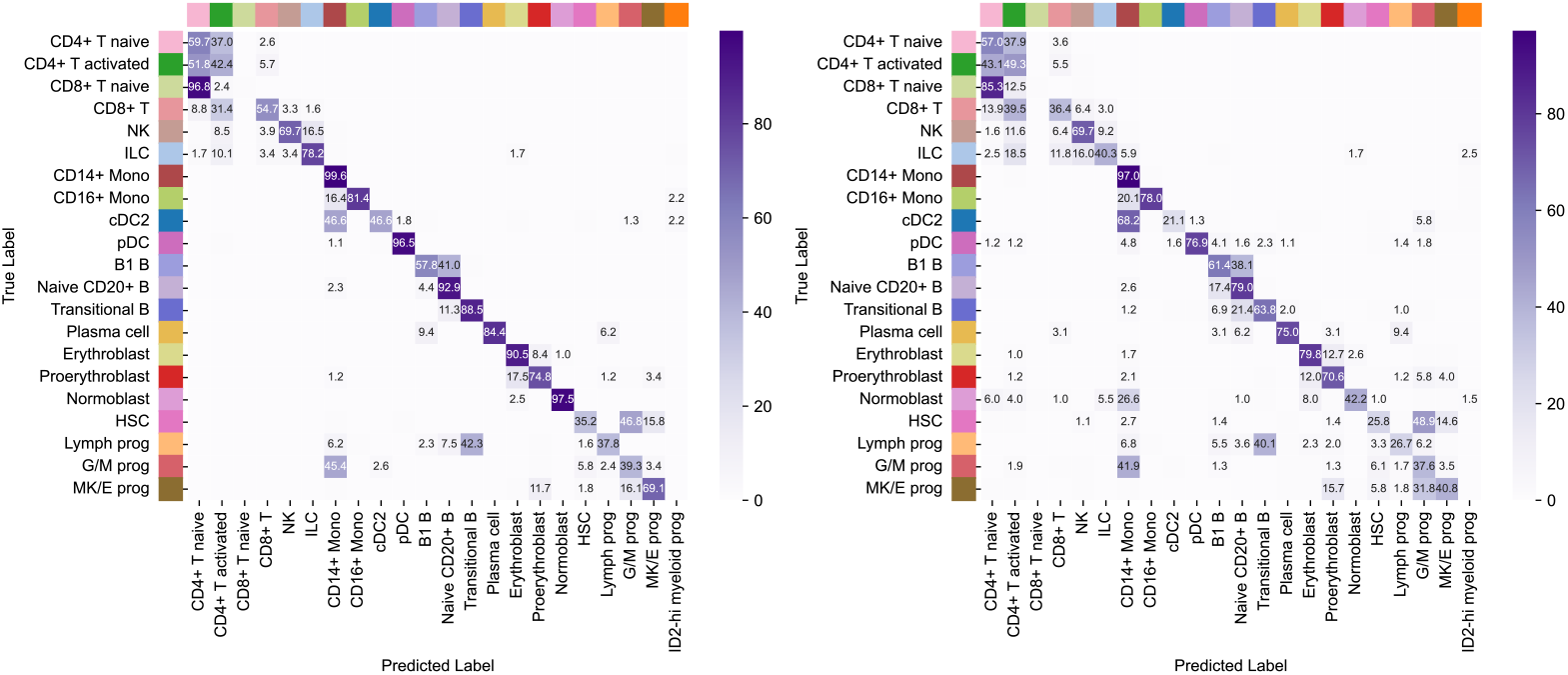
The confusion matrix of the cell type label transfer from the bone marrow single-cell dissociated reference to the full transcriptome (left) and pseudo Xenium panel query set (right).

**Fig. S5:**
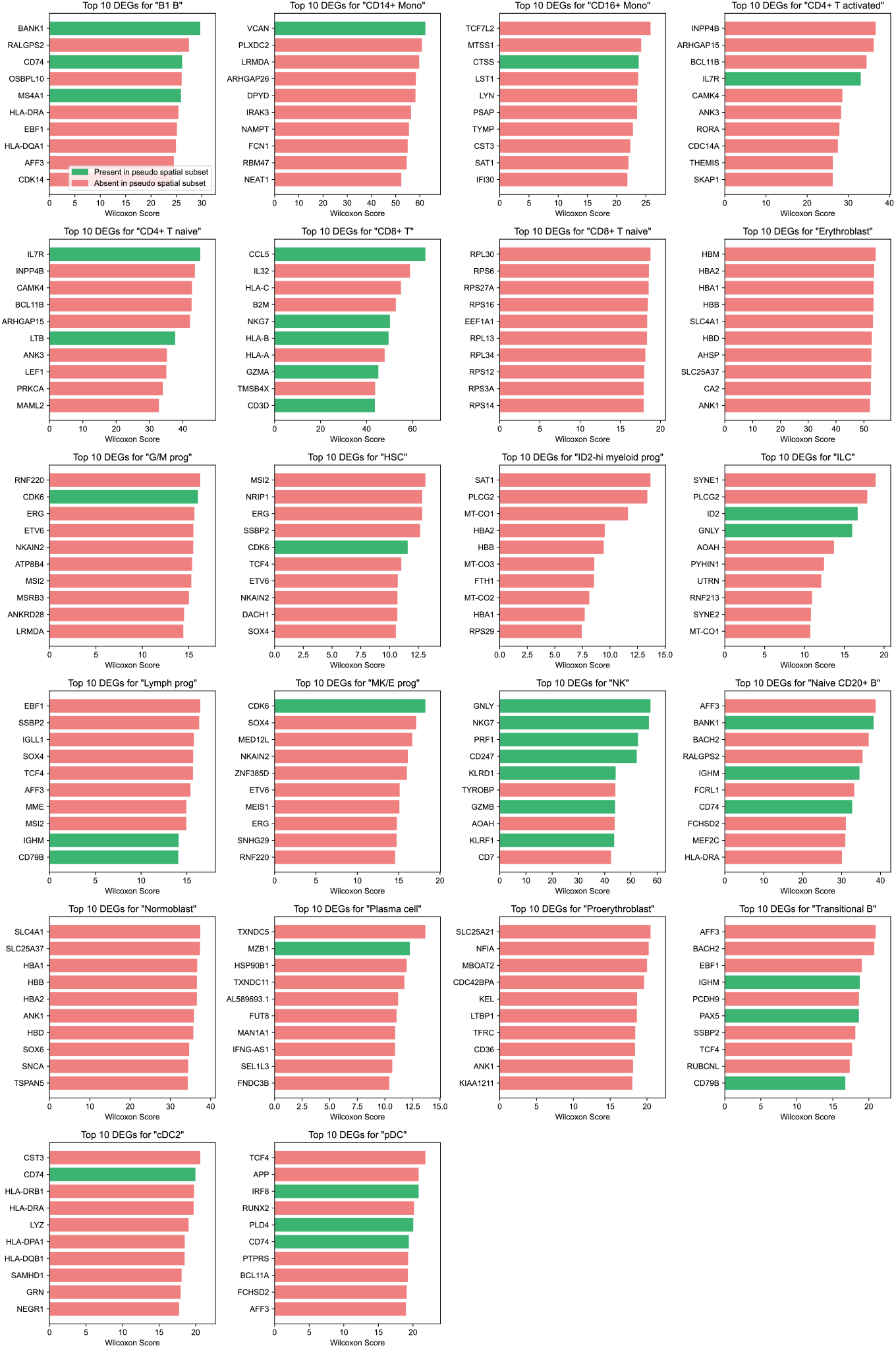
The top differentially expressed genes for each group of cell types compared to the rest of the cells for the bone marrow dataset, sorted by the Wilcoxon score. The genes are colored green if they also exist in the Xenium panel by which the pseudo-spatial data is created, and therefore green if they are dropped in the Xenium panel.

**Fig. S6:**
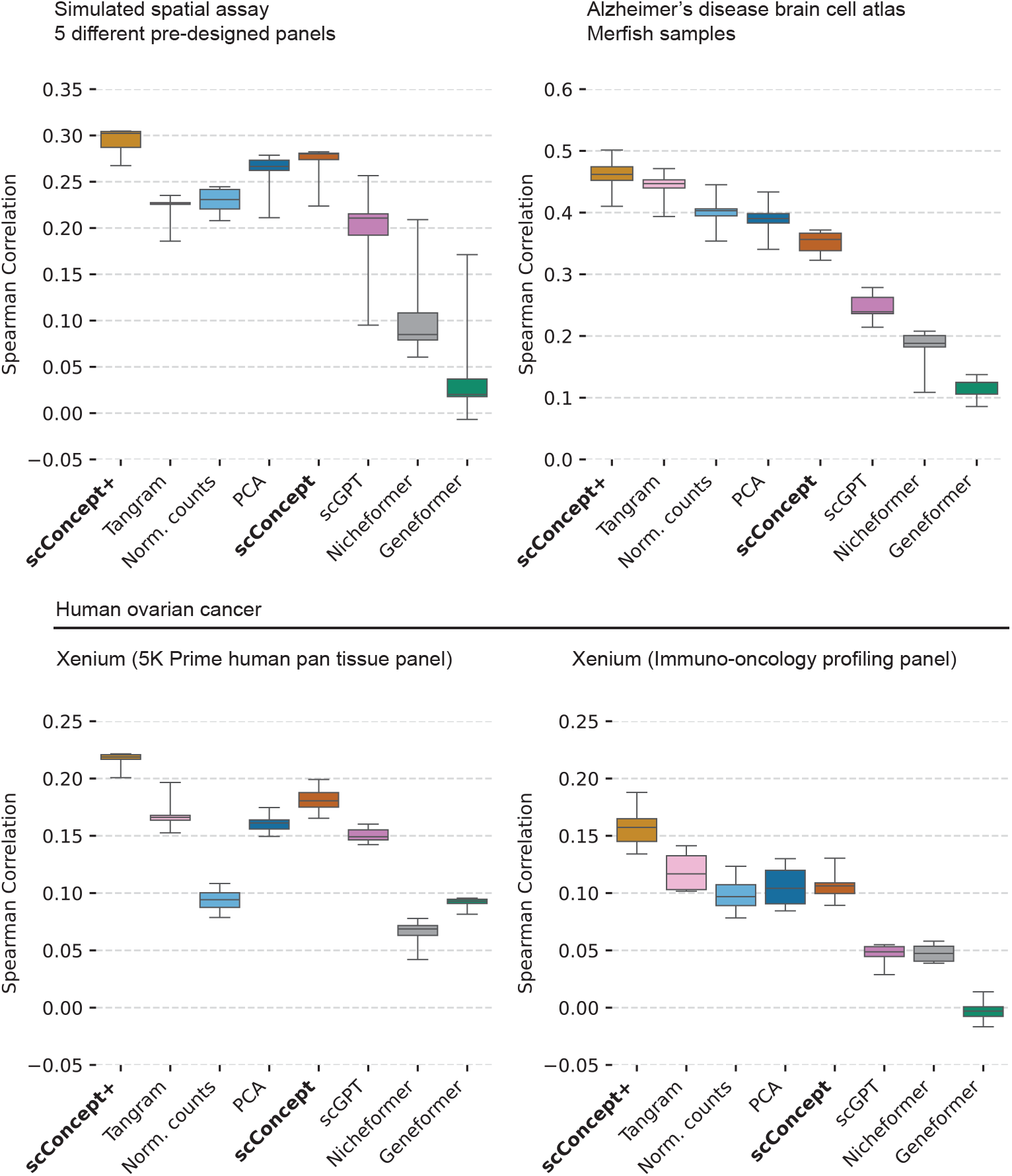
The Spearman rank correlation coefficient evaluation of the imputation performance across methods and datasets.

**Fig. S7:**
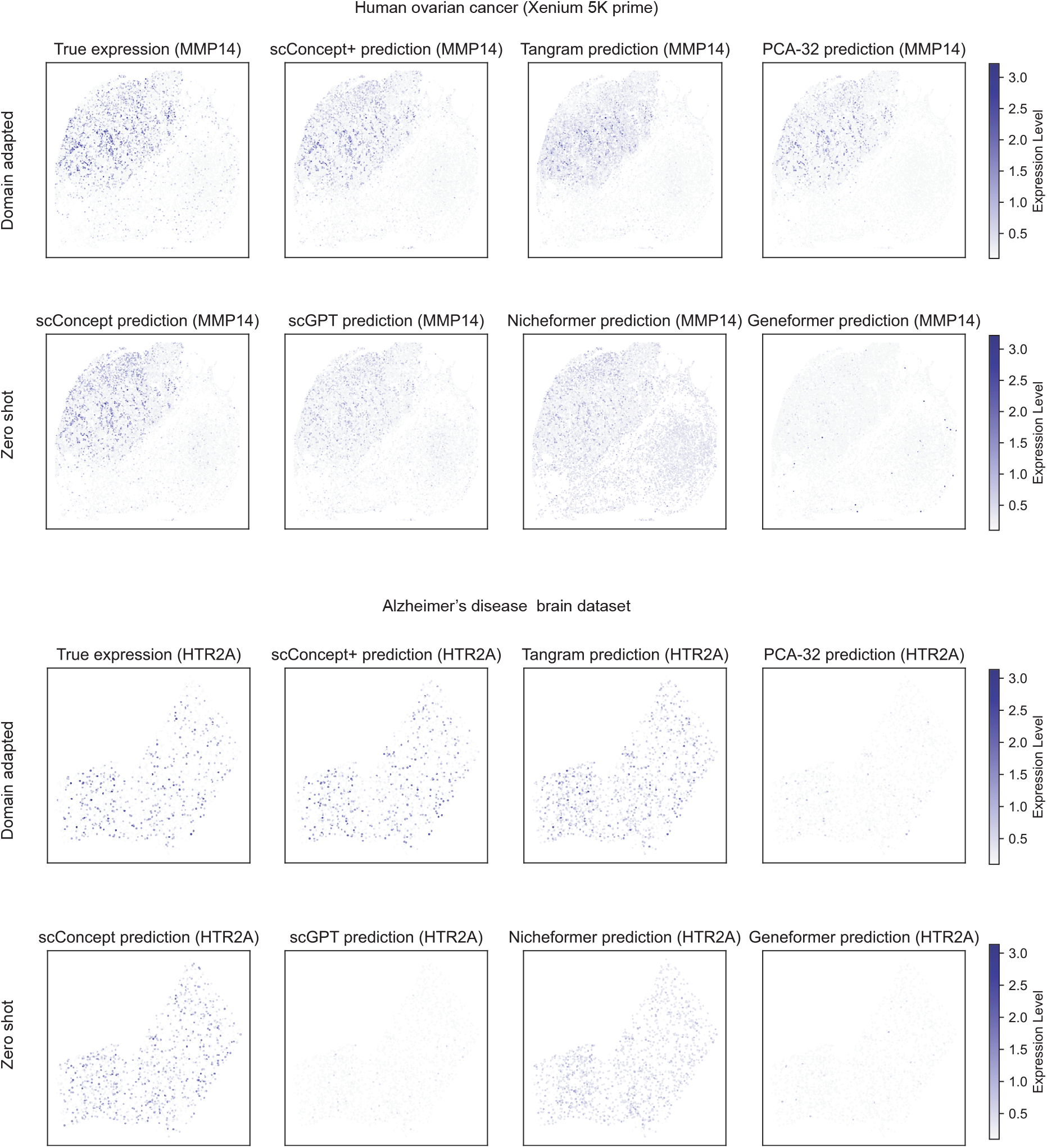
The spatial distribution of the true expressions and the imputed values for the gene MMP14 of the Xenium ovarian cancer dataset and HTR2A of the MERFISH Alzheimer’s disease brain dataset across different methods.

**Fig. S8:**
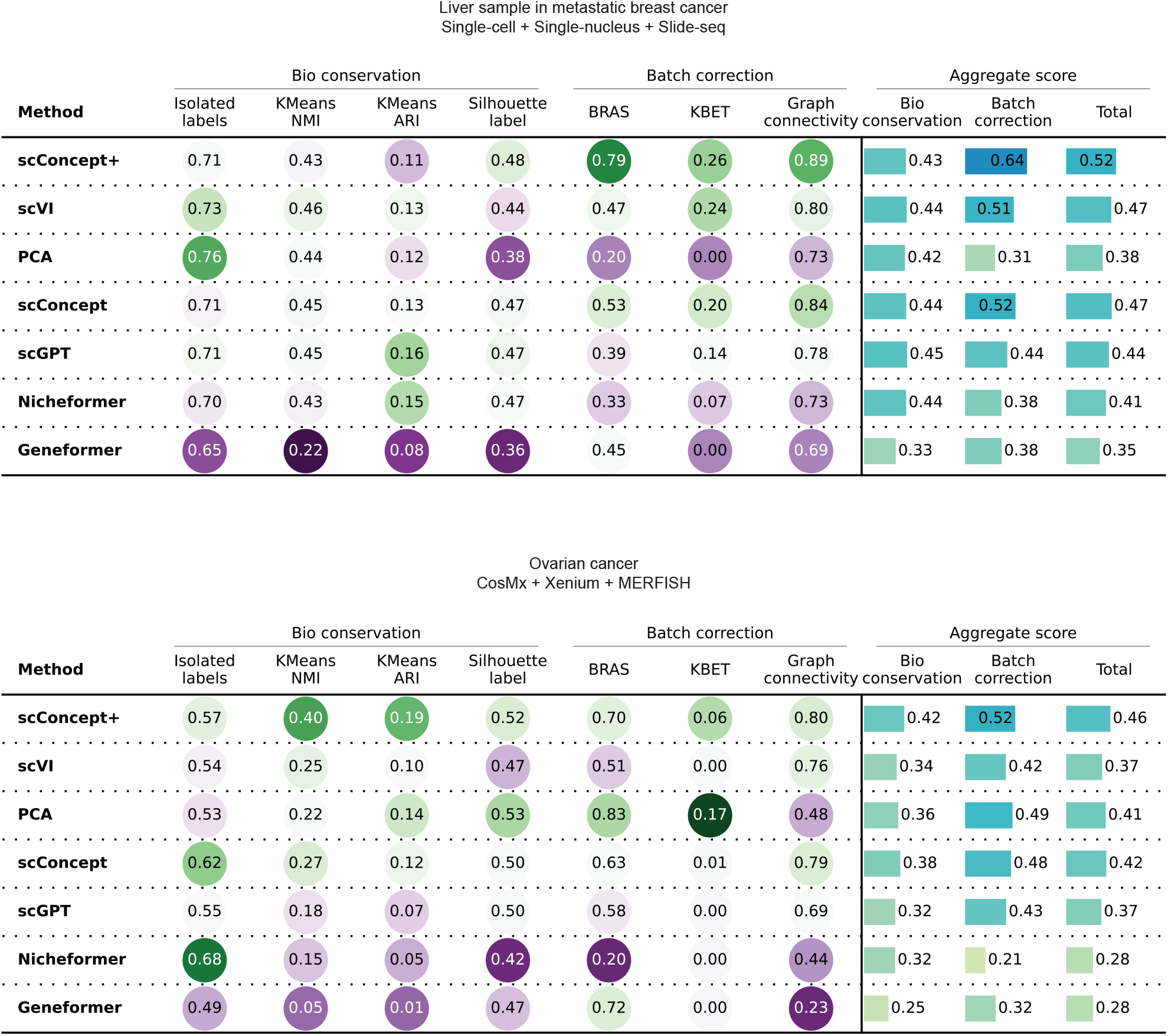
Benchmarking the robustness of cell embeddings through evaluating the data integration of different single-cell assays: 1. Integration of single-cell, single-nucleus, and slide-seq assays of liver samples in metastatic breast cancer. 2. The integration of CosMx, MERFISH, and Xenium spatial assays from human ovarian cancer with different gene panels.

